# Heterogeneous constraint and adaptation across the malaria parasite life cycle

**DOI:** 10.1101/2025.02.11.636054

**Authors:** Sarah A. Perkins, Daniel E. Neafsey, Angela M. Early

## Abstract

Evolutionary forces vary across genomes, creating disparities in how traits evolve. In organisms with complex life cycles, it is unclear how intrinsic differences among discrete life stages impact evolution. We looked for life history-driven changes in patterns of adaptation in *Plasmodium falciparum*, a malaria-causing parasite with a multi-stage life cycle. Categorizing genes based on their expression in different life stages, we compared patterns of between- and within-species polymorphism across stages by estimating nonsynonymous to synonymous substitution rate ratios (*dN/dS*) and mean pairwise nucleotide diversity (𝞹_NS_/𝞹_S_). Considering these alongside estimates of Tajima’s *D*, fixation probability, adaptive divergence proportion and rate, and *F_ST_*, we looked for changes in the drift-selection balance in life stages subject to transmission bottlenecks and changes in ploidy. We observed signals of reduced selection efficacy in genes exclusively expressed in sporozoites, the parasite form transmitted from mosquitoes to humans and often targeted by vaccines and monoclonal antibodies. We discuss implications for how parasites evolve to resist therapeutics and consider functional, molecular, and population genetic factors that could contribute to these patterns.

## Introduction

Within a species, evolution does not uniformly impact all regions of the genome. Mutation, selection, recombination, drift, and gene flow may occur at different rates across loci dependent on factors ranging from chromatin accessibility^1^ to tissue-specific expression.^2^ Differences in selection strength between coding and non-coding regions result in lower substitution rates in the former, underscoring pervasive negative selection against primarily deleterious change in protein sequences impacting organismal fitness. Strong positive selection may shift allele frequencies in only a subset of targeted genes.^3^ Factors like ploidy and effective population size (N_e_) can also intrinsically differ across an organism’s genome. For example, adaptive substitutions accumulate more quickly in genes on sex chromosomes because the effects of recessive variants in these genes are not always masked by heterozygosity.^4^ In this way, molecular, cellular, and organismal characteristics shape evolutionary patterns across genomes.

The life history concept refers to major biological transitions in the life cycle of an organism, such as maturation and reproduction, which are vital to the organism’s evolutionary fitness. Life history traits including fecundity and life span have been demonstrated to explain levels of genetic diversity across phylogenetically diverse organisms.^5^ Within-species trade-offs between life history traits are not uncommon; complex life cycles with morphologically distinct forms are hypothesized to aid adaptation at discrete life stages by decoupling selected traits,^6^ although our full understanding of decoupling, synergism, and antagonism across life stages is still developing.^7,8^ However, with a few exceptions like diplontic plants^9^ and animal gametes,^10^ less attention has been given to how the intrinsic characteristics of distinct life stages may themselves drive patterns of diversity within a single species’s genome. Particularly in single-celled or non-metazoan organisms, complex life cycles can comprise extreme changes, not only in niche and morphology but also in ploidy and clonal cell count. This suggests that life stages may differ both in the types of selection pressures experienced and in the efficacy of the adaptive response to those pressures.

Among these differences, population size and ploidy are likely to exert strong effects on selection efficacy. Modeling approaches have shown how these effects can arise in haploid-diploid life cycles,^11,12^ in which modeled organisms develop and grow in alternating haploid and diploid stages. In a stochastic genetic model of a two-stage haploid-diploid life cycle where selection only occurs in one stage, Bessho and Otto demonstrated that the fixation probability of adaptive mutations decreases with relative stage-specific reproductive value and corresponding population size.^11^ Using forward genetic simulation to model the complex life cycles of plants, Sorojsrisom *et al.* showed an analogous trend when comparing fixation probabilities between stage-specific adaptive mutations, observing an order of magnitude reduction in the chance of fixation when selection occurs in a stage with a reduced population size.^12^ This builds on evidence that haploid-specific genes evolve faster in numerous haploid-diploid plant species.^9^ These findings have important implications for how selection proceeds in complex life cycles, but would be strengthened by empirical validation in life cycles with more complex features, including haploid-dominance, increased stage number, and more drastic demographic transformations.

Vector-transmitted parasites, like the malaria-causing apicomplexan *Plasmodium falciparum*, are excellent models for empirically studying how selection at distinct life cycle stages impacts genomic diversity. As *P. falciparum* parasites pass between human and mosquito hosts, they experience severe population contractions and expansions,^13^ changes in ploidy, and sophisticated stage-specific alterations in gene regulation^14^ enabling invasion of and survival within diverse tissues (Fig. 1). It is clear that the selection landscape is highly variable across this complex life cycle. However, it is not known if the parasite’s ability to adapt to these selection pressures varies simultaneously.

**Figure 1.**
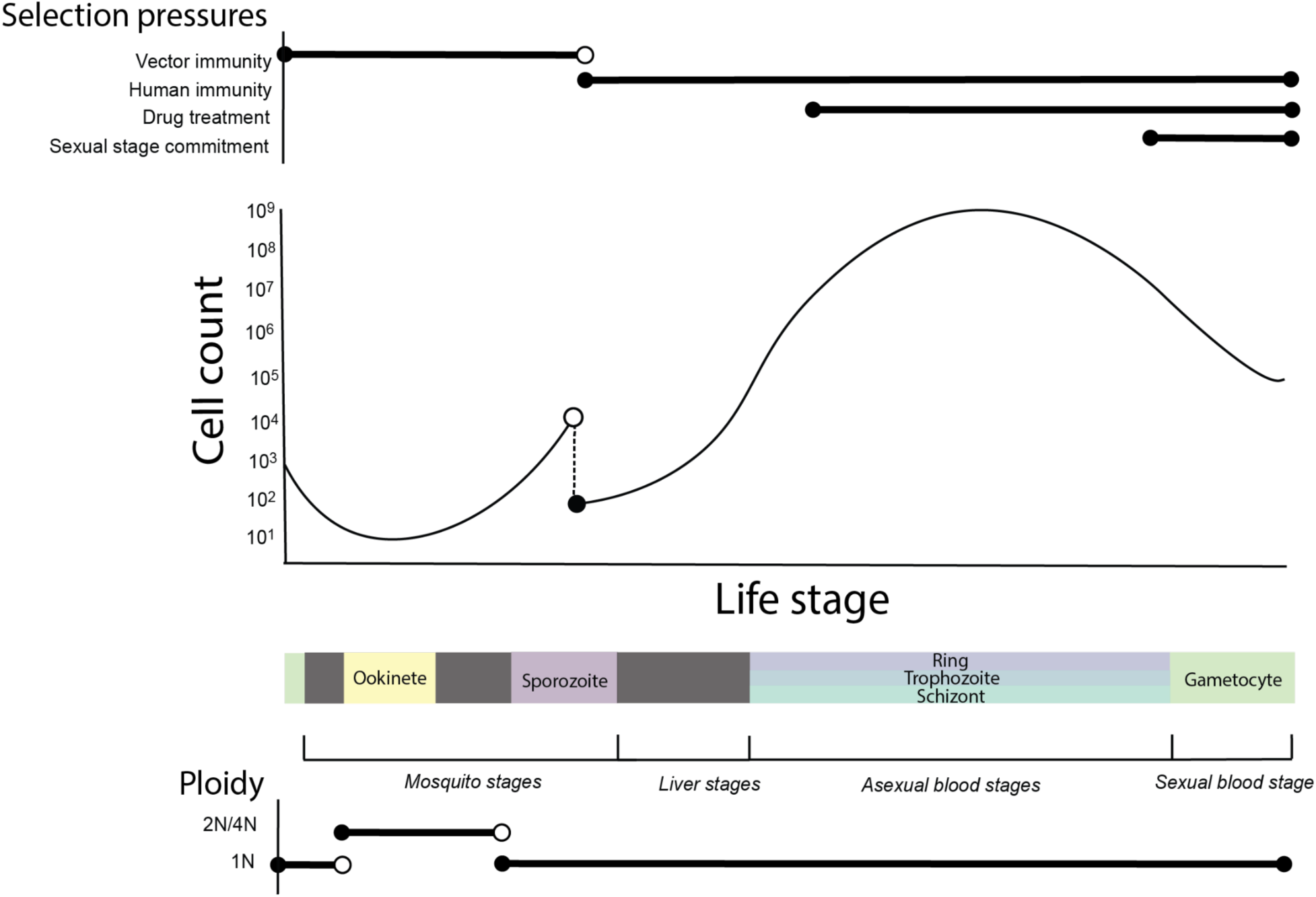
Changes in parasite cell count and ploidy coincide with diverse physiological and therapeutic selective pressures (approximated with horizontal lines) across the infectious life cycle of *P. falciparum*. Colored boxes mark stages used in subsequent analyses and figures. The dashed line denotes a transmission bottleneck.

The *P. falciparum* life cycle begins when an *Anopheles* mosquito ingests around 10^3^-10^4^ male and female gametocytes^15^ (the parasite’s sexual form). These gametocytes rapidly form gametes within the mosquito’s midgut that fuse to produce diploid zygotes. The zygote undergoes meiosis (without cell division) and develops into a tetraploid ookinete that burrows through the mosquito’s midgut epithelium. During the subsequent stage, thousands of haploid nuclear divisions take place within a multinucleated oocyst. Finally, invagination of the oocyst occurs to produce nonreplicative, mononucleated sporozoites that migrate to the mosquito’s salivary glands.^16^ During its next blood meal, the mosquito will inject 10-100 of these stored sporozoites into the host’s skin.^17^ Within hours, a subset of the sporozoites successfully transit to and invade the liver, where they asexually multiply 10^4^ - 10^5^ times before exiting into the bloodstream as merozoites.^18^ Each merozoite invades a red blood cell and goes through a round of asexual amplification lasting around 48 hours,^15^ transitioning through the ring, sporozoite, and schizont stages before bursting as up to 36 free merozoites that go on to invade new red blood cells.^19^ During the asexual blood stages, *P. falciparum* cell counts are capable of expanding to around 10^8^-10^12^ causing disease in the human host.^15^ At each asexual replicative round, a small subset of parasites instead commit to sexual development and differentiate into non-replicating male and female gametocytes, which are transmitted back to the definitive mosquito host for sexual reproduction.

At the whole genome level, demographic shifts associated with the *Plasmodium* life cycle are predicted to enhance both genetic drift and selection.^20^ Within the genome, broader expression of genes across the life cycle correlates with greater evolutionary constraint in rodent and primate *Plasmodium* species.^21^ The *Plasmodium* genome furthermore shows evidence of stage-specific differences in evolutionary constraint.^21^ Understanding whether and how the unique life cycle of *P. falciparum* influences adaptation will not only provide insight into a potentially important evolutionary phenomenon, but also may aid the selection of therapeutic targets–for drugs, vaccines, and monoclonal antibodies–with steeper barriers to resistance evolution.

Here, we test the related hypotheses that selection efficacy and adaptation rate are heterogeneous across the *P. falciparum* life cycle. Using a single cell sequencing dataset from the Malaria Cell Atlas,^22^ we identify sets of genes expressed in single stages. We assess polymorphism in these genes using genomic variation data from four *P. falciparum* parasite populations in the MalariaGen Pf7 database,^23^ quantifying mean gene-level nonsynonymous (𝞹_NS_) and synonymous (𝞹_S_) pairwise diversity, their ratio (𝞹_NS_/𝞹_S_), Tajima’s *D*,^24^ and Hudson’s *F*_ST_.^25^ We also examine between-species nonsynonymous (*dN*) and synonymous (*dS*) substitution rates and their ratio (*dN/dS*) across the genome, comparing *P. falciparum* genes to orthologs in closely related species. We find that, as in other *Plasmodium* species, more broadly expressed genes are more constrained. Further, we consider genes expressed only in single stages and observe signals of enhanced genetic drift in the sporozoite stage, indicating an excess accumulation of non-adaptive alleles. We use DFE-alpha^26,27^ to examine selection efficacy and adaptation rates in each stage, estimating that the sporozoite stage has the lowest selection efficacy among the stages considered. We argue that it is important to extend investigations of stage-specific adaptive potential to inform the development and deployment of parasite therapeutics.

## Methods

### Identification of life stage specific gene sets

To identify the life cycle stages at which each *P. falciparum* gene is expressed, we performed differential gene expression analysis on the Malaria Cell Atlas *P. falciparum* Smart-seq2 dataset.^22^ This dataset leverages the high per-cell coverage of Smart-seq2 single cell sequencing technology^28^ to capture gene expression in parasite cells across six life stages.

Some life stages contained smaller subcategories (for example, salivary gland sporozoite and injected sporozoite); however, to create gene sets with sufficient sizes for this analysis, we utilized the low resolution life stage annotations: ookinete, sporozoite, ring, trophozoite, schizont, and gametocyte.

We identified variable genes through dropout-based feature selection on normalized counts with the `M3DropFeatureSelection` function from M3Drop^29^ (v1.30.0), using an FDR threshold of 0.01. This function selects genes with higher dropout rates than expected in the focal Smart- seq2 assay given their expression level.^29^ From the resulting set of highly variable genes, we defined genes that were expressed in only a single stage via the `M3DropGetMarkers` function to determine ‘marker’ genes with significantly upregulated expression in single cell types, which we considered evidence for cell type-specific expression. To validate patterns of polymorphism in these gene sets, we also develop alternative gene sets with single-stage expression using previously described approaches^14,21^ and a sporozoite gold standard gene set^30^ (Supplementary Appendix).

We excluded expected targets of immune-mediated balancing selection from both gene sets. As a proxy for antigenicity, we used reported serum antibody reactivity from a microarray-based screen of samples from a malaria-exposed cohort.^31^ We removed any protein with a domain recognized by serum antibodies—producing an intensity signal at least 1 SD above the control standard—in at least 10% of the cohort.^31^ We used ShinyGO v. 0.81^32^ to perform a gene set enrichment analysis for the resulting primary gene sets relative to the genomic background.

### Characterization of breadth of gene expression across the life cycle

For each gene, we calculated expression breadth as the number of life stages in which it was expressed, using the same six stages as the stage-specific analysis: ookinete, sporozoite, ring, trophozoite, schizont, and gametocyte. To generate binary *gene x stage* expression classifications based on Malaria Cell Atlas scRNAseq data, we normalized the raw *P. falciparum* Smart-seq2 data set, discarding cells with transcript counts outside the range of 150-2500.

Subsequently, we set a minimum threshold for the proportion of gene-expressing cells per life cycle stage needed to classify a gene as expressed in that stage, defined as 50% of the total number of cells based on prior analyses.^14,21^ We examined the Pearson correlation between the resulting binary gene expression profiles of different stages, including genes expressed in multiple stages. Subsequently, to reduce noise in overall breadth estimates from stochastic variation in background transcript detection–which could falsely increment breadth, particularly through signals of expression in smaller cell populations–we required that genes identified by this procedure demonstrate significant differential expression, via `M3DropGetMarkers`, *uniquely* in the grouped stages they were identified in compared to all other stages, grouped.

This comparison was only performed for genes with breadth < 6, such that assay cells could be grouped into two groups for differential expression testing. Based on these classifications, we generated non-overlapping gene sets grouped by overall breadth of expression of each gene across the life cycle, where breadth values denote the maximum total number of life stages in which a gene was expressed by these criteria. A subset of genes could not be classified as expressed in any stages by these criteria, demonstrating limitations of this filtering approach.

### Population-level characterization of diversity and drift

To characterize population-wide patterns of polymorphism across life stage-specific gene sets, we used whole-genome variant calls from the MalariaGEN Pf7 data set.^23^ We used data from four countries: Ghana, Tanzania, the Democratic Republic of the Congo, and Cambodia, which we chose on the basis of adequate sample size (N >= 75 after filtration) and expected variation in transmission level and genome-wide parasite diversity.

To remove comparisons between highly related parasites, we calculated relatedness (as identity-by-descent; IBD^33^) between all monogenomic (Fws > 0.95) parasites within the same country. We formed clusters of individuals at a threshold of IBD>=0.25 with the R package iGraph and retained only one sample—the sample with the highest proportion of called sites— for downstream analyses.^34,35^ After filtering, the sample sizes from Ghana, Tanzania, the Democratic Republic of the Congo, and Cambodia were 362, 136, 152, and 75, respectively. We only retained biallelic single nucleotide polymorphisms (SNPs) that passed quality control and had VQSLOD values above 2. In addition, on a per-sample basis, we only retained calls that were homozygous and had minimum read depths of 5. We used SNPEff and SNPSift (v. 4.1) to annotate and subset synonymous (S), nonsynonymous (NS), and fourfold degenerate (FFD) variants from the filtered VCF files.

For each gene across the *P. falciparum* 3D7 genome, we calculated the number and proportion of NS, S, and FFD sites. We multiplied resulting counts by the variant call quality filter pass rate to adjust for missing sites (Supplementary Appendix, Fig. S1, Supplementary file 1). Using these corrected counts, we calculated gene-level NS (𝞹_NS_) and S (𝞹_S_) pairwise diversity, as well as their ratio (𝞹_NS_/𝞹_S_), and Tajima’s *D*.^24^ We expect demographic effects to manifest genome- wide and not be impacted by stage of expression, and therefore assume that between-stage differences in diversity are caused by factors attributable to distinct life cycle stages. Finally, divergence between populations was examined through estimation of Hudson’s *F*_ST_ of SNPs segregating in compared populations. We calculated these measurements with scikit-allel.^36^

To examine genome-wide correlations of diversity statistics with gene coding length, we used Kendall’s 𝝉 -B, a rank-based test, on untransformed data. For genes with 𝞹_NS_ > 0 and 𝞹_S_ = 0 or *dN* > 0 and *dS = 0*, we included 𝞹_NS_/𝞹_S_ and *dN/dS* estimates as infinite values. To test for stage-level differences in diversity in each population, we used linear regression to test the association of categorical stages with gene-level diversity statistics, correcting for gene coding length. Prior to regression, we log-transformed coding length and the statistics *dN*, *dS*, *dN/dS*, 𝞹_NS_, 𝞹_S_, and 𝞹_NS_/𝞹_S_, but not Tajima’s *D* and Hudson’s *F_ST_*, to accommodate expected log-linear relationships. In parallel with previous analyses,^37^ to avoid excluding zeros in log-transformed diversity statistics, we incremented all estimates by constant values set based on the minimum observed non-zero estimates of each within each population sample. We also replaced negative *F_ST_* estimates with zeros. We used Tukey tests on resulting models to test for differences in mean statistic values by stage, correcting comparisons for multiple hypothesis testing via the Benjamini-Hochberg procedure. Considering the resulting p-values together, we estimated the significance of meta-population differences in pairwise stage comparisons of diversity statistics via adaptively weighted Fisher (AW-Fisher) tests for combining association statistics.^38^ Finally, to test for changes in 𝞹_NS_/𝞹_S_ and *dN/dS* with breadth, we estimated partial Spearman’s rank correlation coefficients,^39^ controlling log-transformed estimates for log-scaled gene coding length.

### Estimation of ω, α, and Q

We generated NS, S, and FFD folded site frequency spectra for each life stage for all four examined populations via scikit-allel. Resulting spectra were reformatted for input into the DFE- alpha^26,27^ `est_dfe` executable as described in DFE-alpha documentation.

We estimated sequence divergence between *P. falciparum* and both *P. praefalciparum* (divergence time ∼50 kya) and *P. reichenowi* (divergence time ∼190 kya)^40^ using ortholog group classifications from PlasmoDB.^41–43^ We downloaded orthologous pairs of coding sequences from the *P. falciparum* 3D7,^44,45^ *P. praefalciparum* G01,^40^ and *P. reichenowi* CDC^46^ reference genomes for all genes in our life stage-specific gene sets. We aligned translated sequences in Biopython via the pairwise2 `globalms` alignment function (with the following custom scoring parameters: 5 points for matches, -2 points for mismatches, -3 for opening a gap, and -0.5 for extending a gap), kept the best alignments by score for each unique *P. falciparum* gene in the ortholog set, and filtered out alignments with scores below 3.5. Across each aligned sequence pair, we counted NS, S, and FFD nucleotide variants at each codon, excluding gaps and indels. From available high quality alignments, we obtained divergence for 3,643 *P. falciparum-P. reichenowi* and 3,652 *P. falciparum*-*P. praefalciparum* gene pairs. We used these estimates to generate divergence files for input into the DFE-alpha `est_alpha_omega` executable. To generate outlier-robust estimates of average adaptive divergence proportion in each life cycle stage, we iteratively applied DFE-alpha to subsets of each gene set reduced via a leave-one-out approach. The resulting jackknifed estimates of the probability of fixation of a deleterious mutation (*Q*), proportion of adaptive substitution (*α*), and relative rate of adaptive divergence (*ω*) were compared pairwise via the nonparametric Wilcoxon rank sum test.

We use a two-epoch DFE-alpha model correcting for a population size expansion from 10 to 1000 in an optimized generation time (Supplementary Appendix). Although we focus on results using this model with S neutral sites, the reference species *P. reichenowi*, and the reference parasite population sampled in Ghana, we also test alternative analysis parameters (a one-epoch model, FFD neutral sites, the reference species *P. praefalciparum*, and the reference populations sampled in DRC or Tanzania).

## Results

### Single-cell sequencing captures variability in P. falciparum gene expression across six life cycle stages

We used single-cell Smart-seq2 *P. falciparum* data from the Malaria Cell Atlas project^22^ to examine expression patterns across six *P. falciparum* life cycle stages (ookinete, sporozoite, ring, trophozoite, schizont, and gametocyte). The *P. falciparum* reference genome contains 5,280 genes,^44^ of which a subset are reliably analyzed with short-read sequence data and considered part of the “core” genome.^47^ 4,930 of these core genes were detectably expressed in at least 2.5% of one cell type in the assay. We excluded around 20% of detected genes with low (≤70%) variant call pass rates from population diversity statistic calculations (Fig. S1A). In our evolutionary analysis, we also wished to exclude genes subject to strong immune selection. We classified 405 of the expressed core genes (8.2%) as potential antigens based on a large serum antibody reactivity screen^31^ and excluded these from the main analyses. This left us with a set of 3,699 expressed nonantigenic genes for downstream characterization in parasite genomic variation data.

We classified genes by whether or not they were expressed in over 50% of the cells of each of the six life cycle stages. Examining the Pearson correlation of these binary profiles across the stages, we found negative correlations between stages in different host environments, indicating some compartmentalization of the transcriptome by stage physiology (Fig. S2, ring-ookinete *r* = -0.213, *p* < 0.001; ring-gametocyte *r* = -0.178, *p <* 0.001; schizont-ookinete *r* = -0.0748, *p* < 0.01; trophozoite-ookinete *r* = -0.127, *p* < 0.001). From these profiles, we characterized the breadth of expression of genes across the six discrete life cycle stages, further applying a differential expression filter (Methods). Amongst genes that could be classified, we estimated that most non-antigenic genes were detected in only a single stage, and very few genes were expressed across the full life cycle (*N* = 900, 402, 145, 51, 24, 6 for breadth categories 1-6, Supplementary file 1).

Next, we applied a more robust feature selection procedure to isolate genes expressed in only one life stage. We identified non-antigenic genes that were highly variable at the assay level, with significantly increased expression in a single stage. We further excluded genes with breadth estimates above 1. The resulting gene sets ranged in size from 16-54 genes (Fig. S3, Supplementary File 2), with the fewest genes in the trophozoite category and the most genes in the gametocyte category. This finding is consistent with the latter stage encompassing morphologically distinct male and female forms. We repeated this procedure without the feature selection step to define alternative gene sets with single stage expression (Fig. S3, Supplementary File 3).

### P. falciparum nucleotide divergence and diversity correlate with gene length

Gene length may correlate with evolutionary rate directly or indirectly via phylogenetic depth, intronic burden,^48^ alternative splicing, or expression level.^48–50^ *P. falciparum* has unusually long genes and intergenic regions compared to other unicellular eukaryotes and protists, with a high proportion of genes over 4 kb.^51,52^ We examined the effect of gene coding length genome-wide in *P. falciparum*, considering both signals of diversity and divergence. We found that *dN*, *dS*, and *dN/dS* correlate positively with coding length (Fig. S4A-C, Kendall’s 𝝉 = 0.347, 0.279, 0.197; *p <* 0.001*, p <* 0.001*, p <* 0.001). The proportion of NS sites increased with coding length, while the proportion of S and, correspondingly, FFD sites declined (Fig. S4D-F, Kendall’s 𝝉 = 0.260, - 0.260, -0.264; *p <* 0.001*, p* <0.001*, p* < 0.001). Using signals of within-species diversity recovered from 3,699 *P. falciparum* genes in a deeply sampled African population (Ghana; *N*=362), we found that coding length positively correlates with 𝞹_NS_ and 𝞹_S_ and negatively correlates with 𝞹_NS_/𝞹_S_ and Tajima’s *D* (Fig. S5, Kendall’s 𝝉 = 0.190, 0.264, -0.0978, -0.605; *p* < 0.001, *p* < 0.001, *p <* 0.001, *p* < 0.001).

Transcript length can also influence the sensitivity of detection in RNA sequencing assays. Accordingly, we find that coding length correlates with breadth (Kendall’s 𝝉 = 0.200, *p <* 0.001). We also note that coding length varies in our single-stage gene sets (Fig. S6C-D). Although this could represent a meaningful biological phenomenon, we aimed to test for changes in evolutionary rate independent of coding length to exclude assay-level biases as drivers of our observations. In subsequent analyses, we include coding length as a covariate to correct for this effect.

### Increase in selective constraint with breadth of expression across the life cycle

We aimed to determine whether expression breadth correlates with signatures of adaptive constraint in *P. falciparum.* Based on a previous analysis of *dN/dS* in rodent *Plasmodium* spp. by Tebben *et al.*,^21^ we hypothesized that greater breadth of expression across the parasite life cycle would be associated with signatures of higher selective constraint, such that more broadly expressed genes would be more highly conserved both between and within species compared to more narrowly expressed genes.

We calculated a covariate-adjusted Spearman’s rank correlation coefficient that controlled for log-scaled gene coding length to analyze the genome-wide change in *P. falciparum-P. reichenowi dN/dS* with expression breadth, and we found a small but statistically significant negative correlation (Fig. S7A, Partial Spearman’s *𝝆* = -0.112, *p* = 1.32 x 10^-4^). We found the same negative association of breadth with *dN/dS* estimates from *P. falciparum-P. praefalciparum* sequence comparisons (Fig. S7B, Partial Spearman’s *𝝆* = -0.0969, *p* = 0.00122).

Next, we test a parallel comparison of expression breadth with population-level diversity. We found a weak negative correlation of expression breadth with 𝞹_NS_/𝞹_S_ in three of the four populations, but the pattern is not statistically significant (Fig S7C). We also find an enrichment of nuclear functions amongst genes expressed in more than three stages (Fig. S8).

### Genes expressed in single life stages do not show distinct functional enrichments

We next examined functional enrichment of the single-stage gene sets (Fig. S8). These sets were limited to genes expressed in only one stage. We hypothesized that different functional enrichments could indicate functional divergence between non-overlapping gene sets, and additionally wished to determine whether any large skews existed which in and of themselves could explain divergent evolutionary patterns across life stages. We found primarily mixed enrichment signals, including enrichments of uncharacterized labels in the sporozoite, trophozoite, and ookinete single-stage gene sets. We found no functional enrichments in the gametocyte single-stage gene set. In the schizont single-stage gene set, we uncovered some defined functional enrichments, including: myristate, phosphoprotein, ANK repeat, lipid catabolic process, repeat, and coiled coil. These hits may indicate a slight skew toward essential functions like lipid catabolism and toward protein-protein interactions, consistent with the intensive developmental shifts and growth occurring during this stage. However, these labels may also point to more thorough structural characterization of gene products in this set. While gaps in functional annotation across the *P. falciparum* genome^44^ and the modest size of gene sets limit functional inference, we do not find clear evidence for strong functional differences between single-stage gene sets outside of a slight skew toward some metabolic and protein-protein interaction functions in the schizont set.

### Substitution patterns vary across life stages

We next tested whether signatures of species-level divergence show variation across discrete life cycle stages. To do so, we used log-linear regression to compare *dN*, *dS*, and *dN/dS* for *P. falciparum*-*P. reichenowi* ortholog pairs encompassing genes expressed in only one *P. falciparum* life cycle stage. We observed significantly higher *dN* and *dN/dS* in sporozoite-limited genes compared to genes expressed in only one of the other five stages (Fig. 2A, *dN* comparisons: *Q < 0.01, 3/5; Q < 0.001, 2/5.* Fig. 2B, *dN/dS* comparisons*: Q < 0.05, 3/5; Q < 0.01, 1/5; Q < 0.001, 1/5)*. Genes with expression limited to the *P. falciparum* sporozoite stage had a median *dN*/*dS* value of 0.411, over twice that of all other genes with single-stage expression (median *dN/dS* = 0.184). We also observed elevations in gametocyte *dN* and *dN/dS* (Q < 0.05, both comparisons) compared to the schizont stage (Fig. 2A). We saw no evidence that the life cycles varied in basal mutation rate, as *dS* levels were comparable across all gene sets (Fig. 2C). Rather, the higher sporozoite-stage *dN/dS* appears to be driven by *dN* increases (Fig. 2A). These results are consistent with work by Tebben *et al.* who found higher mean *dN/dS* values in genes expressed in the sporozoite stage of the rodent parasite *P. berghei.*^21^

**Figure 2.**
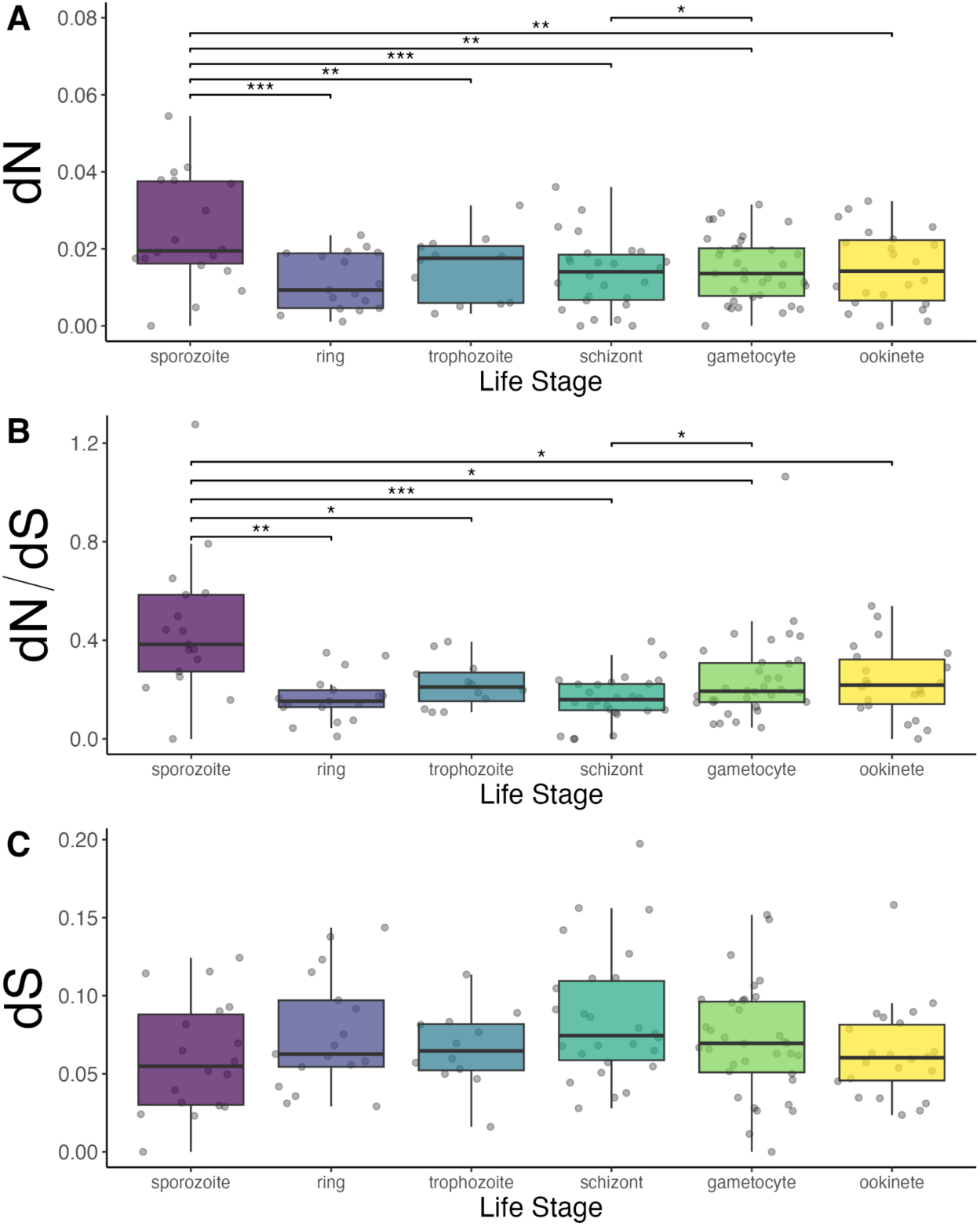
Gene-level nonsynonymous (A) and synonymous (C) *P. falciparum-P. reichenowi* divergence rates and their ratio (B), separated by life stage (x-axis/color). All estimates (A-C) were compared linearly by stage after log-transformation and correction for log-transformed coding length as described (Methods) (A) BH-adjusted significant pairwise dN comparisons: sporozoite-ring, *p* = 5.97 x 10^-4^; sporozoite-trophozoite, *p* = 0.00553; sporozoite-schizont, *p* = 7.95 x 10^-5^; sporozoite-gametocyte, *p* = 0.00844; sporozoite-ookinete, *p* = 0.00553; gametocyte-schizont, *p* = 0.0297. (B) BH-adjusted significant pairwise dN/dS comparisons: sporozoite-ring, *p* = 0.00222; sporozoite-trophozoite, *p* = 0.0419; sporozoite-schizont, *p* = 9.20 x 10^-5^; sporozoite-gametocyte, *p* = 0.0344; sporozoite-ookinete, *p* = 0.0194, gametocyte-schizont, *p* = 0.0144 [1 infinite y-axis value not represented]. (C) All BH-adjusted pairwise dS comparison *p* > 0.05.

### Patterns of polymorphism vary across life stages

To examine patterns of polymorphism across genes expressed in single life cycle stages, we used data from the MalariaGen network documenting parasite variation within four geographically diverse populations.^23^ We calculated gene-level diversity metrics including 𝞹_NS_, 𝞹_S_, 𝞹_NS_/𝞹_S_, and Tajima’s *D* (Supplementary file 1). Overall, the Tajima’s *D* estimates show a strong negative skew, a well established pattern in *P. falciparum* populations.^20,53,54^

We used linear regression to assess the association between gene-level diversity estimates and categorical life stage, examining pairwise differences in mean estimates for each stage. To aggregate stage-level trends across all four separate populations, we used AW-Fisher p-value meta-analysis on the resulting population-level associations. We found increases in 𝞹_NS_ and 𝞹_NS_/𝞹_S_ for genes only expressed in the sporozoite stage relative to those only expressed in four of the other five stages (Fig. 3A-B, Table S1, *Q* > 0.05 sporozoite-trophozoite 𝞹_NS_ comparison; *Q* < 0.01, sporozoite-ookinete 𝞹_NS_, 𝞹_NS_/𝞹_S_ and sporozoite-trophozoite 𝞹_NS_/𝞹_S_ comparison; *Q* < 0.001, all other comparisons). However, we did not generally see this trend for 𝞹_S_, a proxy for neutral sequence change, although we did observe an elevation of 𝞹_S_ in ookinete genes compared to sporozoite and gametocyte genes (Fig. 3D, Table S1, *Q <* 0.01, sporozoite- ookinete 𝞹_S_ comparison and *Q* < 0.001, gametocyte-ookinete 𝞹_S_ comparison). Genes expressed only in the sporozoite stage also showed higher Tajima’s *D* values than those expressed in four of the five alternative stages, while ookinete genes had higher *D* values than ring stage genes (Fig. 3C, Table S1, *Q* > 0.05, sporozoite-ookinete *D* comparison; *Q* < 0.05, sporozoite- trophozoite *D* comparison and ookinete-ring *D* comparison*; Q* < 0.001, all other comparisons).

**Figure 3.**
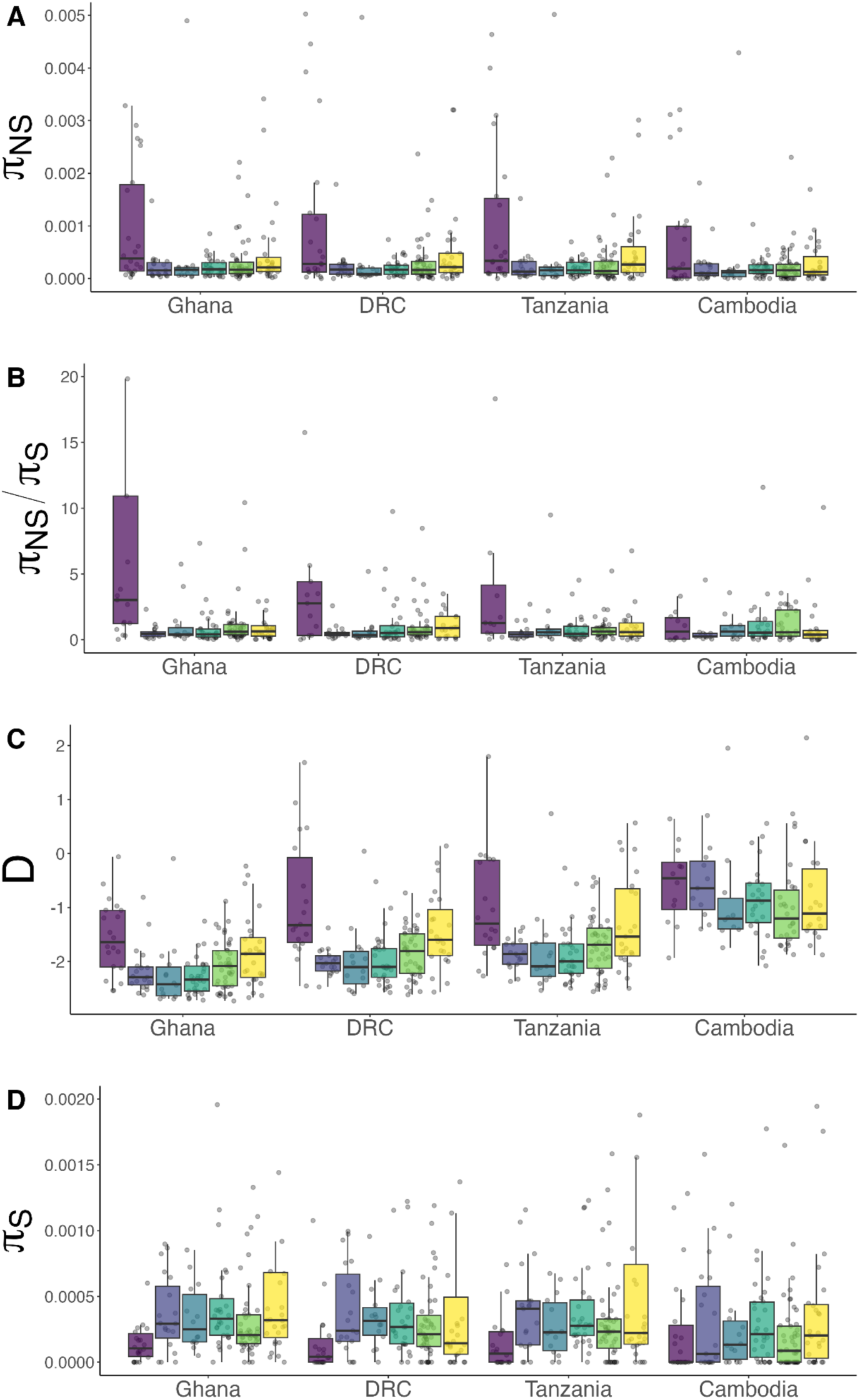
Gene-level population diversity statistics including 𝞹_NS_ [2 outliers above y-axis limit not shown] (A), 𝞹_NS_/𝞹_S_ [9 outliers above y-axis limit not shown and 105 infinite values not represented] (B), Tajima’s *D* (C), and 𝞹_S_ [22 outliers above y-axis limit not shown] (D) across four geographically diverse MalariaGen parasite population samples (x-axis). Estimates are shown for genes with single-stage expression and grouped by life stage, which is denoted by boxplot color (purple: sporozoite, periwinkle: ring, blue: trophozoite, blue-green: schizont, green: gametocyte, yellow: ookinete).

We obtain comparable results when comparing these estimates between our alternatively defined gene sets with single stage expression (Fig. S9, Table S2).

### Population-level genetic differentiation varies across life stages

Next, we compared genetic polymorphism across parasite populations. We hypothesized that population differentiation may differ by stage of expression, which could be attributed to either geographically variable selection on stage-specific functions or reduced constraint (higher drift) operating at some life cycle stages. We estimated per-gene Hudson’s *F_ST_* in pairwise comparisons of four parasite populations sampled in Ghana, Tanzania, the Democratic Republic of the Congo, and Cambodia (Supplementary file 1).

Africa is the inferred point of origin for *P. falciparum* with the parasite likely spreading to Southeast Asia 50 to 60 kya.^55^ We used linear regression to test for pairwise *F_ST_* differences between stages, again aggregating the results of population comparisons with AW-Fisher meta-analysis of p-values. We found that the sporozoite stage had a higher *F*_ST_ than all other stages except the schizont stage (Fig. 4, Table S3, *Q* < 0.05, 2/4 comparisons; *Q* < 0.001, 2/4 comparisons). These differences are most pronounced for the within-Africa comparisons (Fig. 4A).

**Figure 4.**
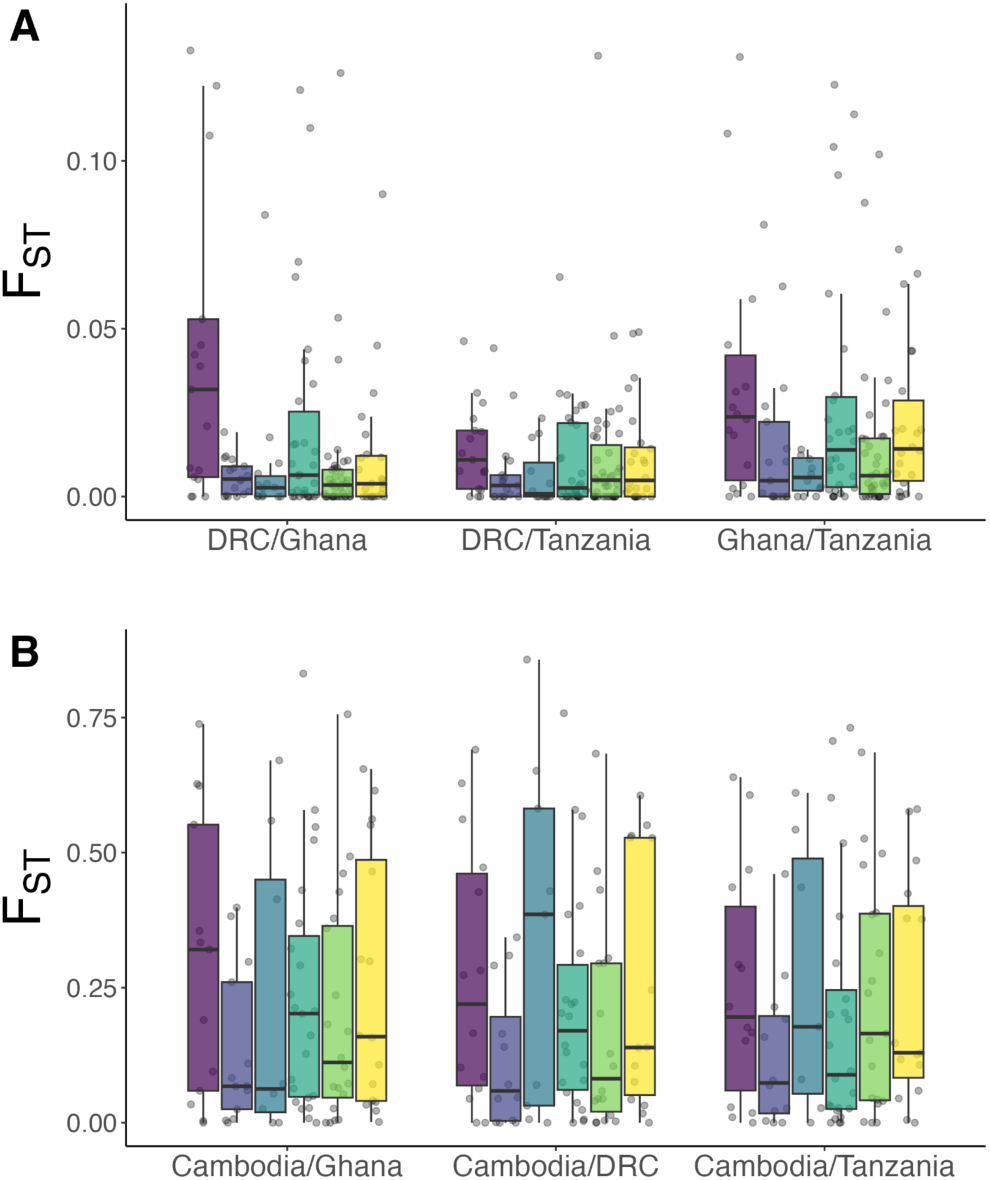
Hudson *F_ST_* estimates comparing *P. falciparum* populations, defined here at the country level, within the African continent (A) and between Africa and Asia (B). [4 outliers above y-axis limit in (A) are not shown]. Life stage is denoted as boxplot color (purple: sporozoite, periwinkle: ring, blue: trophozoite, blue-green: schizont, green: gametocyte, yellow: ookinete).

### Signals of reduced selection efficacy in the sporozoite and ookinete stages

After finding that expression in different life stages may impact how nucleotide substitutions accumulate across the *P. falciparum* genome, we estimated the probability of fixation of novel deleterious mutations and the rate and proportion of beneficial sequence change at the *P. falciparum* species level. We did this separately for each single-stage gene set to test if these fundamental quantities, which reflect the efficacy of natural selection and rate of adaptation, differ substantially between genes according to their functional life history context.

We used DFE-alpha^26,27^ to estimate the fraction of beneficial (adaptive) between-species substitution. Assuming that most within-species polymorphism is deleterious, DFE-alpha uses site frequency spectra generated from population data to estimate the distribution of fitness effects (DFE) for each gene set and generate expectations for the proportion of effectively neutral (slightly deleterious) mutations. These expectations are compared with observed levels of non-neutral (NS) relative to neutral (S or FFD) between-species divergence; excess observed non-neutral divergence is interpreted as adaptive.^26,27^ This analysis produces estimates of the probability of fixation of a novel deleterious mutation (*Q*), the proportion of adaptive divergence (*α*), and the relative rate of adaptive substitution (*ω*) for each gene set.

Relying on a jackknifing approach to generate outlier-robust comparisons of gene sets with single-stage expression, we assessed *Q*, *α*, and *ω* for each gene set category (Fig. 5). These estimates were all significantly different from each other across stages (*p* < 0.001, all comparisons). We show an increase in *Q* for the sporozoite stage (Fig. 5A, median *Q*_sporozoite_ = 9.88 x 10^-4^) relative to all other stages and particularly the blood stages (median *Q*_non-sporozoite_ = 2.15 x 10^-4^, median *Q*_ring | trophozoite | schizont_ = 8.46 x 10^-5^). These estimates imply that a novel deleterious mutation is over ten times more likely to fix if it arises in a gene expressed only in the sporozoite stage compared to genes expressed only in one of the blood stages. Genes expressed only in gametocytes or ookinetes are estimated to fix deleterious polymorphism over twice as often as genes expressed only in an asexual blood stage (median *Q*_ookinete_ = 2.31 x 10^-4^, median *Q*_gametocyte_ = 2.16 x 10^-4^).

**Figure 5.**
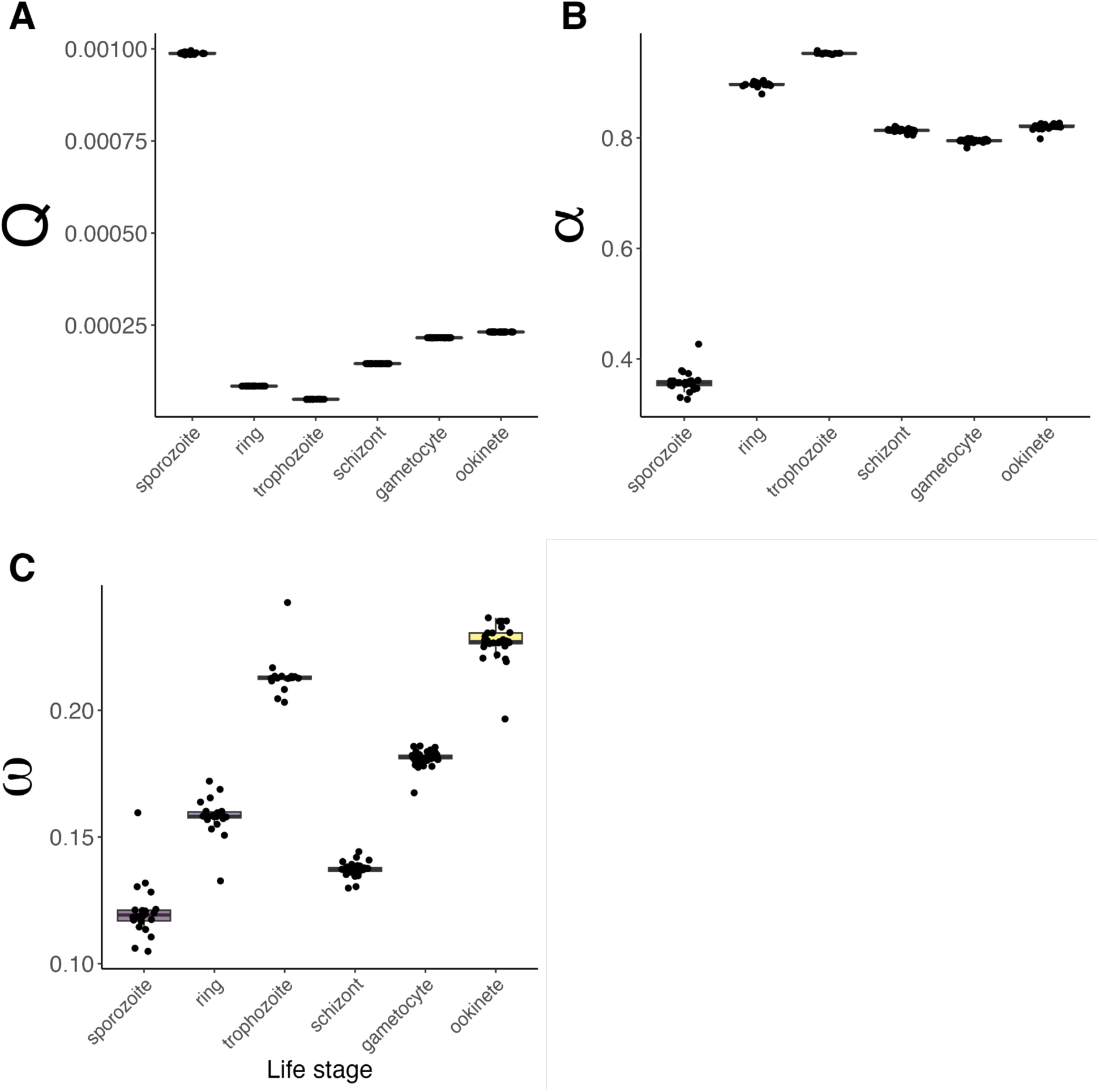
Jackknifed estimates of probability of fixation of a deleterious mutation (*Q*), Proportion of adaptive substitution (*α*), and the relative rate of adaptive divergence (*ω*) estimated by DFE- alpha analysis of site frequency spectra and *P. falciparum*-*P. reichenowi* divergence estimates for nonsynonymous (selected) and synonymous (neutral) SNPs in *P. falciparum* life stage-associated genes. All stage-stage pairwise comparisons of jackknifed estimates revealed significant differences by life stage (Bonferroni-adjusted Wilcoxon rank sum test *p* < 0.001).

Overall, the sporozoite gene set has proportionally fewer adaptive polymorphisms than the other gene sets (Fig. 5B, median *α*_sporozoite_ = 0.357, median *α*_non-sporozoite_ = 0.814). We obtain similar relative elevations in Q and reductions in α for the sporozoite gene set when we use FFD sites rather than S sites as neutral sites (Fig. S10A, C, E), when we analyze *P. falciparum-P. praefalciparum* divergence (Fig. S10B, D, F), or when we use population data from the other African parasite populations (Fig. S11A-D).

### Directional selection peaks in the trophozoite and ookinete stages

Unlike *dN/dS*, which does not distinguish between adaptive and non-adaptive coding changes, our *ω* estimates are adjusted for differences in selection efficacy to reveal underlying differences in directional selection. The sporozoite stage gene set had the lowest estimated rate of adaptive divergence (*ω*), while the ookinete and trophozoite sets had the highest (Fig. 5C).

The ring and schizont stages also have low *ω* values in the range of those estimated for sporozoites in some jackknife iterations (Fig. 5C). Considering *Q* and *ω* jointly, these results are consistent with a predominance of negative over positive selection in ring and schizont stages, while trophozoite and ookinete stages seem to experience the most positive selection. However, *ω* estimates were more sensitive to changes in analysis parameters and the exclusion of single genes than *Q* and *α* estimates (Figs. 5, S10-S12).

When we only consider substitutions at FFD sites to be neutral (Fig. S10A, C, E) or when we investigate divergence between *P. falciparum* and its closest known relative *P. praefalciparum* rather than *P. reichenowi* (Fig. S10 B, D, F), we observe similar *Q* and *α* rankings, with sporozoite genes maintaining the lowest proportion of adaptive mutations. However, we find differences in the relative *ω* rankings of stages in these iterations, largely driven by increases in sporozoite *ω* estimates (Fig. S10E, F). The median sporozoite *ω* estimates increase 1.5- to 2- fold in these iterations, representing the second highest median *ω* in the latter (Fig. S10E, F, median *ω_sporozoite,_* _S10E_ = 0.179, median *ω_sporozoite, S10F_* = 0.237).

When the DFE estimation step relies on parasite population data from the DRC and Tanzania, the trophozoite *α* estimates remain the highest among the gene sets but decrease by about 8- 11% (Fig. S11 D-E median *α*_trophozoite, DRC_ = 0.869, median *α*_trophozoite, Tanzania_ = 0.842) compared the estimates using population data from Ghana (Fig. 5B, *α*_trophozoite, Ghana_ = 0.954). This analysis also ranks the sporozoite gene set lowest and the trophozoite gene set highest in terms of adaptive divergence, but assigns the sporozoite gene set implausible (negative) *α* and *ω* values, indicating that the model does not accurately adjust for the demography of these populations (Fig. S11C-F).

## Discussion

As the propensity of *P. falciparum* to evolve resistance to therapeutics poses an ongoing threat to malaria elimination efforts,^15^ evolutionary considerations have gained increased prominence in therapeutic development. Vaccine trials evaluate construct diversity,^56,57^ drug studies dissect patterns of *in vitro* resistance evolution,^15,58^ and policy makers weigh the potential for opposing selection pressures with specific drug combinations.^59^ Here, we foreground an additional consideration: variability in adaptive potential across the parasite life cycle may impact long-term therapeutic success, and may be particularly relevant for the selection of targets and the timing of interventions.

Dissecting life stage-specific patterns of adaptation by isolating them within genes expressed in single stages, we find that selection efficacy may be lowest in mosquito and transmission stages and the sporozoite stage in particular. This is consistent with the expectation that low cell count, transmission bottlenecks, and higher ploidy will reduce selection efficacy. If this explanation is correct, the blood stages are poised to respond more effectively to novel selection pressures than the mosquito and transmission stages. An important caveat is that we conduct our analysis and infer parameters relevant to drift and selection in the context of negative selection. Although we expect that reduced selection efficacy should weaken positive selection in tandem with negative selection by heightening the drift barrier, there is less empirical work documenting the distribution of fitness effects of advantageous mutations,^60^ making it difficult to predict the ultimate consequences of elevated drift for adaptation to novel positive selection pressures.

Overall, however, these results demonstrate that patterns of variation diverge across *P. falciparum* life stages, which calls for further investigations of this phenomenon.

We find population genetic signatures of elevated drift in genes that are expressed only in the sporozoite stage, including increased 𝞹_NS_ and 𝞹_NS_/𝞹_S_ (Fig. 3). These signals demonstrate that nonsynonymous mutations accumulate to a greater extent in these genes, showing inheritance patterns consistent with genetic drift or positive selection. We further estimate that deleterious mutations are ten times more likely to become fixed if they arise in genes expressed only in the sporozoite stage compared to genes expressed in one of the asexual blood stages (Fig. 5). Our analysis of α values indicates that, over time, this effect cumulatively decreases the proportion of adaptive sequence changes in sporozoite stage genes by about 45% relative to asexual blood stage genes, which show unusually high proportions of adaptive polymorphism close to 90%. While we implemented a demographic correction in our model to remove upward bias in *α*, these absolute estimates could reflect residual bias stemming from our model’s inability to implement bottlenecks beyond a few thousand generations in the past. Nonetheless, work in other organisms has shown large variation in α estimates, which have ranged from 25-94% in *Drosophila* species alone,^61,62^ based on the estimation techniques used and genomic regions analyzed.^63^ High *α* values have also been observed in haploid pathogens and commensals,^63,64^ suggesting that the ploidy and life history characteristics of these organisms may enhance adaptation. Whether or not inflations beyond this range represent true elevations unique to *P. falciparum* biology or to the focal gene sets is unclear, but could represent an intriguing future research direction in light of the dually intensified drift and selection driven by the *Plasmodium* life cycle.^20^ Importantly, we expect the relative ranks of stage-level α estimates to be robust to any residual demographic bias as this will inflate α estimates similarly across the whole genome. To this end, we observed that upper and lower α rankings were stable when implementing the demographic model compared to the null baseline model of parasite demography (Fig. S12).

We hypothesize that factors like low parasite numbers, bottlenecks imposed during transmission, or lingering bi-allelic protein expression may contribute to drift-dominated evolutionary regimes on sporozoite-expressed genes. Low parasite cell number may directly reduce selection efficacy by decreasing *N_e_* within that life stage, a primary determinant of the magnitude of the selective effect required for selection, rather than drift, to determine the evolutionary fate of an allele. In addition, the parasite undergoes its highest number of mitotic divisions in the asexual blood stages. Mutations impacting fitness in other stages may arise in the asexual blood stages, drift in the absence of selection, and become stochastically fixed during the subsequent human-mosquito transmission bottleneck. Finally, while the parasite undergoes nuclear meiotic division after forming a zygote (Fig. 1), there is some evidence for diploid protein expression continuing through the sporozoite stage.^65^ This increased functional ploidy could mask deleterious alleles and reduce selection efficacy in these stages relative to stages with strictly haploid protein expression.

Overall, these population genetic signals are consistent with there being lower selection efficacy at the sporozoite, and to a lesser extent, ookinete and gametocyte stages. However, we wish to highlight additional hypotheses regarding these diversity patterns. Among these is the possibility that stronger or more diverse selection pressures act on sporozoites as they cross tissues and evade immune responses in both mosquito and human. Higher rates of between-species coding changes could indicate enhanced positive selection at the species level, while increased *F_ST_* could suggest continued geographically dynamic selective gradients. However, in our *ω* estimation, we found that the sporozoite stage gene set did not tend to have the highest inferred rate of adaptive substitution (Fig. 5C, 10E), except when we estimated divergence from sequence comparisons with *P. praefalciparum* orthologs (Fig. S10F) or when we did not implement the demographic model to correct for the recent *P. falciparum* population bottleneck (Fig. S12).

Genetic draft, in which strong selection on certain alleles boosts the frequency of proximal ‘hitchhiker’ alleles in the same genetic background,^66^ could compound signals of heightened nonsynonymous diversity in the setting of stronger positive selection. The effects of draft are largely indistinguishable from those of drift,^67^ making this explanation difficult to rule out.

However, the high recombination rate of *P. falciparum*^68^ would be expected to weaken this effect by causing decay in linkage disequilibrium between polymorphic sites with nucleotide distance. Given that we observe the population genetic signals across genes on multiple chromosomes, and that no single gene seems to underlie sporozoite-specific elevations in *Q*, *α*, and *ω*, we might reason that this scenario would imply a pervasive elevation in positive selection across genes at this stage.

Across the *P. falciparum* genome, we find evidence that negative selection is stronger when genes are expressed in more stages of the parasite life cycle. We observe modest decreases in *dN*/*dS* with expression breadth, consistent with increased constraint and paralleling an analogous *dN*/*dS* trend in rodent *Plasmodium* species.^21^ We also see a weak negative but not significant decrease in 𝞹_NS_/𝞹_S_ with breadth in the largest parasite population. As stated by Tebben *et al.*, these trends could reflect the tendency of core functional genes to be broadly expressed across the parasite life cycle.^14,21,69^ The upper three breadth categories (4-6) in our analysis were enriched for nuclear functions relative to the genomic background, which is consistent with this hypothesis (Fig. S8). Higher breadth genes may also have increased overall expression, which is known to intensify negative selection against protein misfolding and explain large proportions of variation in *dN/dS* in other single-celled eukaryotes.^37^ Finally, antagonistic pleiotropy across stages can also constrain adaptation to any single niche. It could contribute to the observed decrease in *dN/dS* with breadth, depending on its prevalence across the genome. Across the whole transcriptome, we found signals of inverse regulation of gene expression in more physiologically dissimilar stages (Fig. S2). We speculate that these negative correlations reflect some compartmentalization of transcriptional architecture across the life cycle, consistent with the hypothesis that complex life cycles enable adaptation to distinct selective environments. This architecture likely reduces antagonistic pleiotropy across the genome.

The implications of our observations for different therapeutic selective pressures—such as both positive and negative selection imposed by a novel drug compared with diversifying or balancing selection imposed by a vaccine—warrant further investigation. Simulation studies, building off of those modeling the whole malaria life cycle^20^ or modeling evolution across haploid-diploid life cycles^12^ may be extended to identify specific drivers and predict outcomes of life stage-specific differences in organismic evolution.

## Acknowledgements

This study was supported with federal funds from the National Institute of Allergy and Infectious Diseases, National Institutes of Health, Department of Health and Human Services, under Grant Numbers T32AI049928 and T32AI007535 to the Harvard T.H. Chan School of Public Health and Grant Number U19AI110818 to the Broad Institute. We thank Jacob Tennessen, Philipp Schwabl, Raphael Brosula, and Stephen Schaffner for feedback on various analyses in this manuscript. This publication uses MalariaGEN data as described in ‘Pf7: an open dataset of Plasmodium falciparum genome variation in 20,000 worldwide samples’ MalariaGEN et al, Wellcome Open Research 2023, 8:22 https://doi.org/10.12688/wellcomeopenres.18681.1. This publication also uses Malaria Cell Atlas Project data as described in ‘Single-cell RNA-seq reveals hidden transcriptional variation in malaria parasites’ Reid, A.J. et al, eLife 2018, 7 e33105 https://doi.org/10.7554/eLife.33105. This publication uses data from PlasmoDB, a resource from VEuPathDB, as described in ‘VEuPathDB: the eukaryotic pathogen, vector and host bioinformatics resource center in 2023.’ Alvarez-Jarreta, J. et al., Nucleic Acids Res. 2024, 52, D808–D816 https://doi.org/10.1093/nar/gkad1003. This publication also uses supplementary data described in ‘Novel serologic biomarkers provide accurate estimates of recent Plasmodium falciparum exposure for individuals and communities.’ Helb, D.A. et al., Proceedings of the National Academy of Sciences 2024, 112, E4438–E4447 https://doi.org/10.1073/pnas.1501705112 and in ‘Novel insights from the Plasmodium falciparum sporozoite-specific proteome by probabilistic integration of 26 studies’ Meerstein-Kessel, L. et al., PLOS Computational Biology 2021, 17, e1008067 https://doi.org/10.1371/journal.pcbi.1008067.

## Contributor roles

AME: Conceptualization, Supervision, Methodology, Formal Analysis, Writing- review & editing; DEN: Funding Acquisition, Supervision, Resources, Writing- review & editing; SAP: Data curation, Formal Analysis, Investigation, Visualization, Writing-original draft.

## Competing interests

None.

## Notes

https://github.com/sperkibody/PfalLifeCycleSelection

### Competing Interest Statement

The authors have declared no competing interest.

