## Supplementary material for "Heterogeneous constraint and adaptation across the malaria parasite life cycle": README.txt

Supplementary file 1, diversity_values.csv: output of diversity statistic analysis pipeline. 


Supplementary file 2, m3_combined.csv: life stage labels for primary gene sets. 


Supplementary file 3, stage_specific.csv: life stage labels for secondary gene sets. Labels include sporozoite stage gold standard labels from Meerstein-Kessel, L. et al.
