## Supplementary Appendix for "Heterogeneous constraint and adaptation across the malaria parasite life cycle"

### Supplementary Methods

#### *Correcting gene-level site counts for missing variant data*

As the MalariaGen Pf7 VCF included variant sites only,[1] we did not know if excluded sites were invariant or incapable of being called. Accurate gene-level nucleotide diversity calculations require knowing the total number of called sites within a gene, and so for each gene, we calculated the proportion of called SNPs that passed the Pf7 quality control thresholds. We filtered the dataset to exclude genes below a pass rate of 0.70, which excluded roughly 20% of genes in the genome from the analysis (Fig. S1A, lower bound value=0.70, denoted by black line). This excluded 25 genes from the primary gene sets, leaving 159 total. There was a genome-wide correlation between the total coding length of genes and the calculated pass rate among SNPs in coding regions called in Pf7 (Fig. S1B, Kendall's $\tau = 0.161$, $p < 2.2 \times 10^{-16}$). We did not detect a significant impact of life stage on SNP call pass rate among non-antigens in our primary gene sets post-filtering (Fig. S1C, Kruskal-Wallis $\chi$-squared = 3.47, $p = 0.628$). However, we did detect a significant impact in our secondary gene sets (Fig. S1D, Kruskal-Wallis $\chi$-squared = 29.249, $p = 2.07 \times 10^{-5}$). Finally, for each gene across the *P. falciparum* 3D7 genome, we calculated the number and proportion of NS, S, and FFD sites and multiplied resulting site counts by our missing data proportion estimates to adjust for missing sites in subsequent analyses (Supplementary file 1).

#### *Assembling alternatively defined gene sets with single-stage expression*

We also employed an alternative strategy described previously[2,3] to define single-stage expression as expression in at least 50% of the cells with a given stage label. In addition, to avoid propagating potential noise in the dataset characteristic of single cell RNAseq assays, we required that this expression be significantly different from that observed in all other stages as determined by pairwise Wilcox tests conducted with `M3DropGetMarkers`.[4] With this approach, we found a larger range of gene set sizes (8-321 genes, Fig. S3, Supplementary File 3). In this case, the sporozoite stage set comprised the smallest number of genes and the gametocyte stage set the largest. The lower number of sporozoite stage genes described with this approach may reflect differential detection bias at the stage level due to cell count differences in the filtered assay (cell counts: 141 sporozoite, 63 ring, 51 trophozoite, 50 schizont, 88 gametocyte, and 112 ookinete), lower sensitivity due to reduced transcript length of sporozoite stage transcripts, or lower sensitivity due to reduced transcript count in sporozoite cells (Fig. S6). We expect that our primary gene sets defined using a feature selection step will be less influenced by stochastic dropout of lowly expressed transcripts, particularly in stages with intrinsically lower mean expression levels (Fig. S6A-B). In addition, we expect that the 50% expression threshold determining the alternative gene sets may exclude some stage-specific genes that show high within-stage variation in expression. For the downstream analysis, we therefore focus on the results from the first classification approach, which produced more evenly sized gene sets. However, patterns of polymorphism across gene sets defined by this alternative approach, augmented with a previously published sporozoite stage gene set,[5] are qualitatively similar (Fig. S9, Table S2).

***Adjusting DFE-alpha model for a recent population bottleneck***

Recent demographic expansions following population bottlenecks can upwardly bias estimates of α.[6] To reduce potential bias stemming from the recent bottleneck in *P. falciparum*,[7] we use the two-epoch DFE-alpha model[6,8] to correct for a change in population size across $t_2$ generations. We set the initial population size to 10 and the present day population size to the maximum allowable value (1000), assuming that *P. falciparum* has a large present-day effective population size compared to its population size at speciation. If we assume that *P. falciparum* completes roughly 2-6 generations per year (based on generation time estimates of 60-180 days)[7] and underwent a previously inferred population bottleneck 4,000-6,000 years ago,[7] the number of generations modeled exceeds the numerical capacity of the DFE-alpha `est-dfe` executable, preventing maximum likelihood optimization. We therefore specified that the model use maximum likelihood optimization to estimate $t_2$ since the most recent population contraction. Although this would underestimate true $t_2$, we expect that this should reduce bias due to recent population contraction. We emphasize the relative differences in our stage-limited $Q$, $\alpha$, and $\omega$ estimates given that we expect all portions of the genome to be similarly impacted by demographic events.

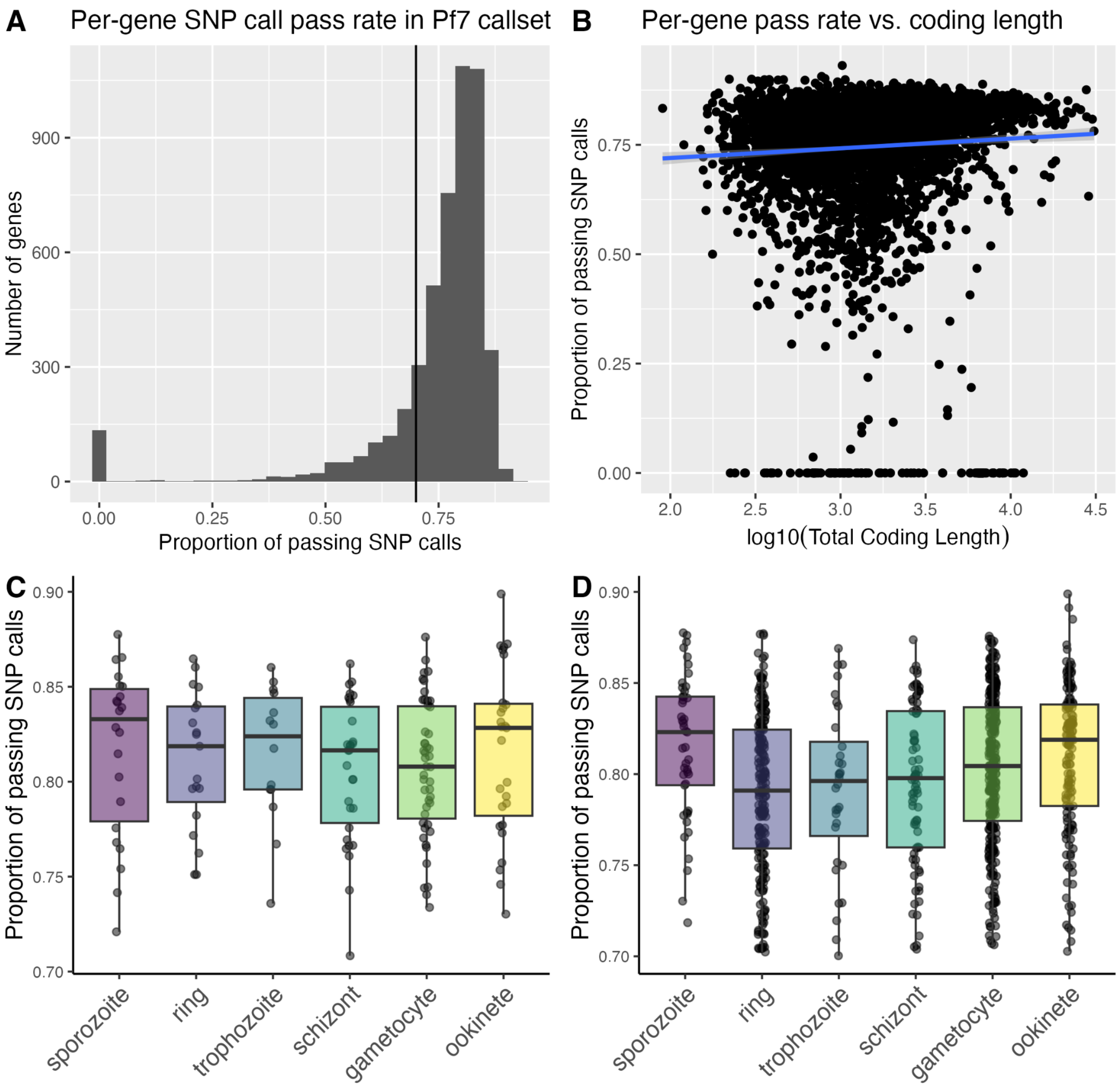


**Figure S1.** Histogram of the proportion of SNP variants in each gene (*N=4925*) that pass variant quality filter checks in the MalariaGen Pf7 variant call data. The proportional variant quality pass threshold used here (0.70) is marked vertically in black (A). Proportional variant quality pass rate per-gene (y-axis) vs. log10-coding length (Kendall's $\tau$ = 0.161, $p < 2.2 \times 10^{-16}$) (B). Proportional variant quality pass rate per-gene (y-axis) vs. life stage labels (x-axis) for the primary gene sets (Kruskal-Wallis $\chi$-squared = 3.47, $p$ = 0.628) (C) and the secondary gene sets (Kruskal-Wallis $\chi$-squared = 29.249, $p = 2.07 \times 10^{-5}$) (D) after excluding antigens and variants below the 0.70 pass rate threshold.

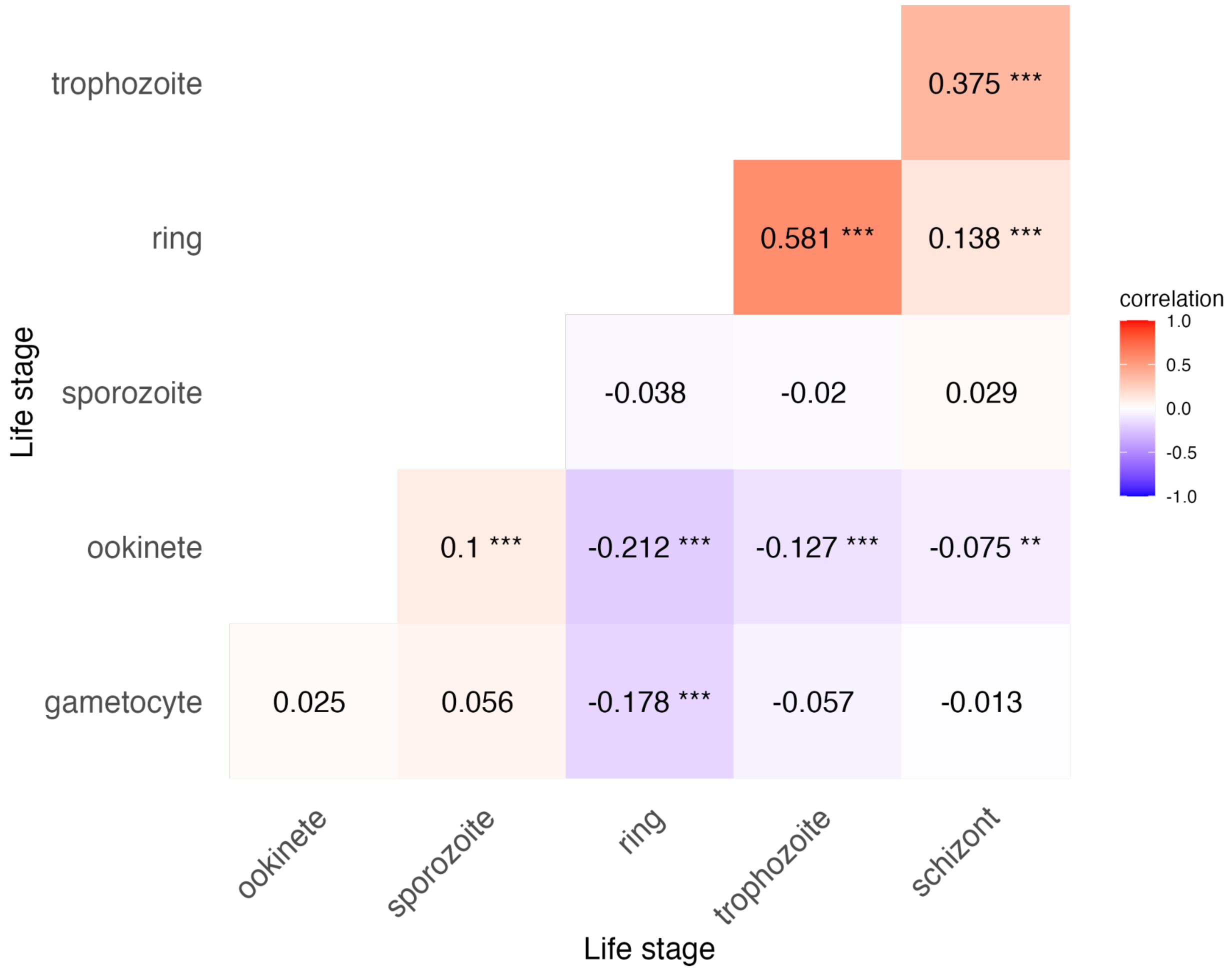


**Figure S2.** The Pearson pairwise correlation of binary gene expression in each stage (color). This calculation includes only non-antigenic genes with detectable expression–i.e., expression in at least 50% of cells–in at least one cell type (stage). Bonferroni-adjusted significant comparisons include: ring-trophozoite $r = 0.581$, $p < 0.001$; ring-schizont = 0.138, $p < 0.001$; schizont-trophozoite $r = 0.375$, p < 0.001, ring-ookinete $r = -0.212$, $p < 0.001$; ring-gametocyte $r = -0.178$, $p < 0.001$; schizont-ookinete $r = -0.0748$, $p < 0.01$; trophozoite-ookinete $r = -0.127$, $p < 0.001$, sporozoite-ookinete $r = 0.0996$, $p < 0.001$. Significant comparisons are marked with asterisks (*$p < 0.05$, **$p < 0.01$, ***$p < 0.001$).

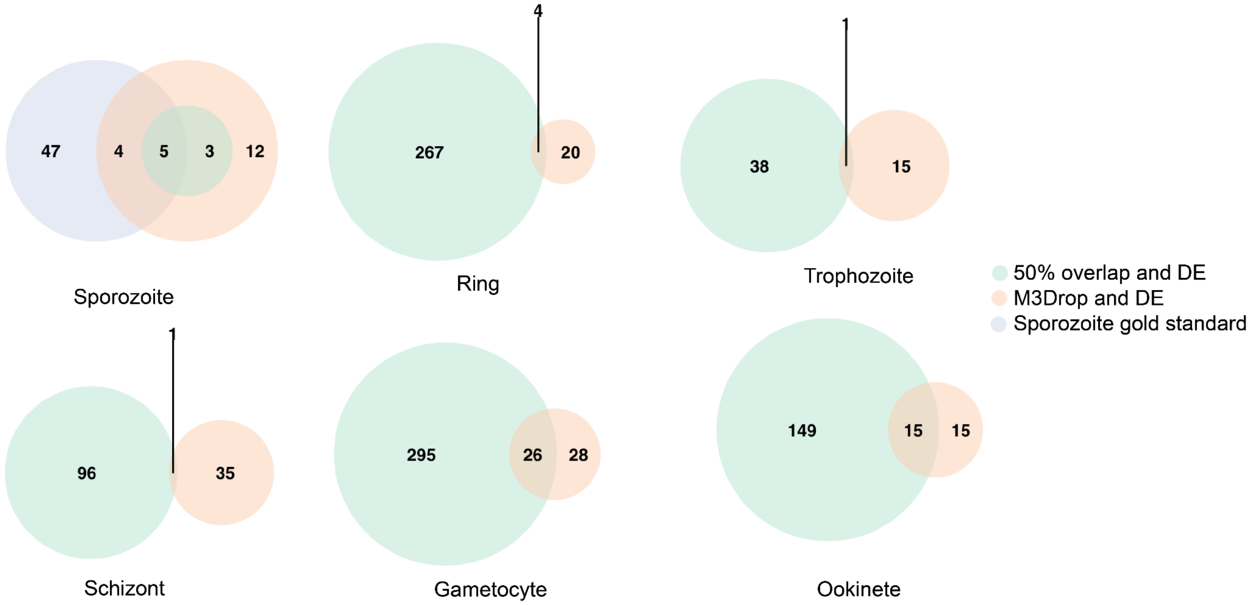


**Figure S3.** Overlap between sets of genes with expression in single stages—displayed here as counts of shared and unshared genes in Venn Diagram format—identified for each low resolution stage in this assay by primary (orange) and secondary (green) gene set assembly methods, after filtering for antigenicity and breadth but prior to combining the sporozoite gold standard gene set (blue, upper left) with the secondary sporozoite gene set (green, upper left).

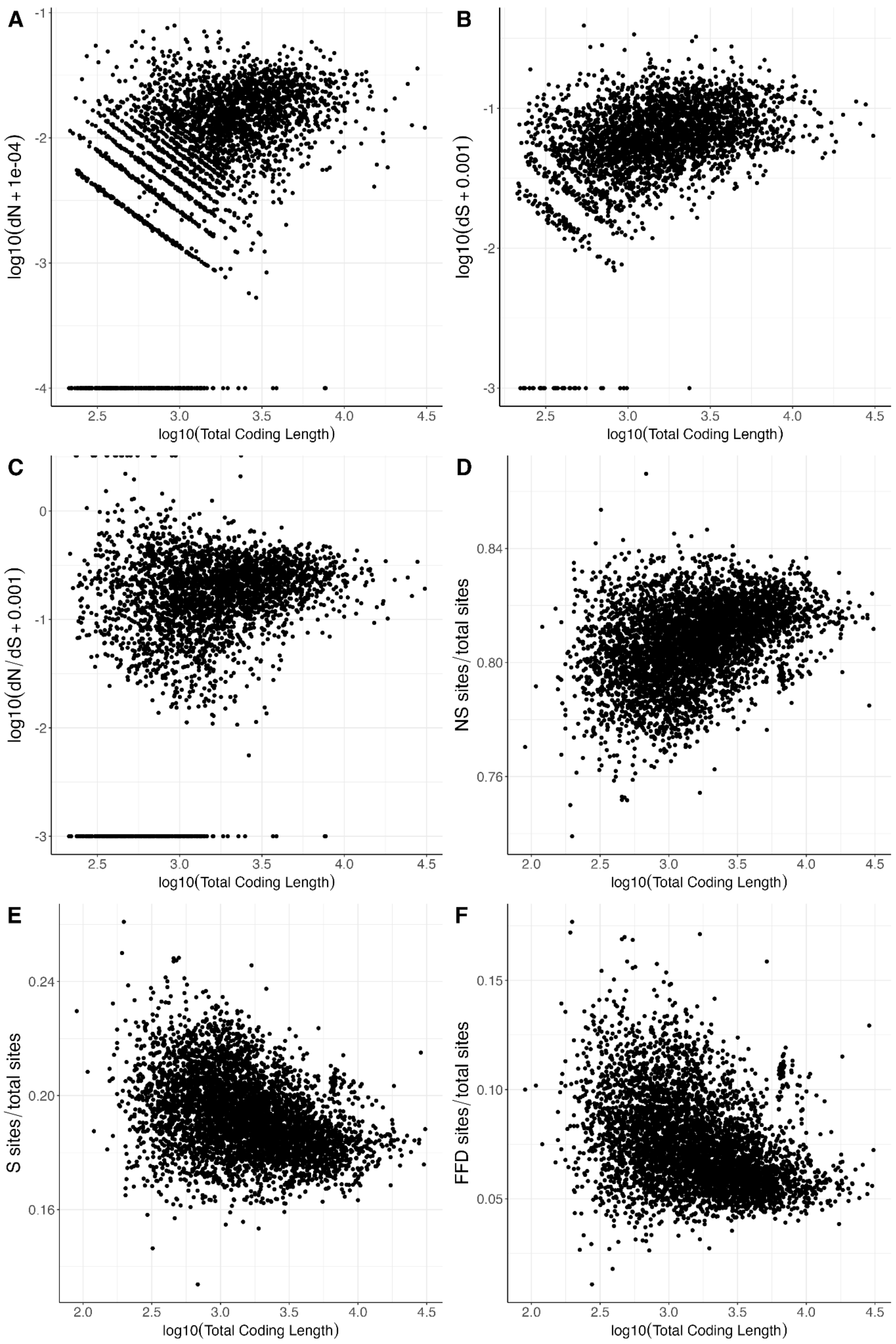
A
log10(dN + 1e-04)
log10(Total Coding Length)
B
log10(dS + 0.001)
log10(Total Coding Length)
C
log10(dN/dS + 0.001)
log10(Total Coding Length)
D
NS sites/total sites
log10(Total Coding Length)
E
S sites/total sites
log10(Total Coding Length)
F
FFD sites/total sites
log10(Total Coding Length)

**Figure S4.** Genome-wide correlations of *dN* (A), *dS* (B), *dN/dS* (C), the proportion of NS sites (D), the proportion of S sites, (E) and the proportion of fourfold degenerate synonymous sites, (F) with the log10-scaled total coding length of transcripts (Kendall's $\tau$ = 0.347, 0.279, 0.197, 0.260, -0.260, -0.264; $p < 2.2 \times 10^{-16}$, $p < 2.2 \times 10^{-16}$, $p < 2.2 \times 10^{-16}$, $p < 2.2 \times 10^{-16}$, $p < 2.2 \times 10^{-16}$, $p < 2.2 \times 10^{-16}$). To avoid excluding zeros in the visualizations, we incremented all values by the lowest observed order of magnitude for the statistic (A-C)

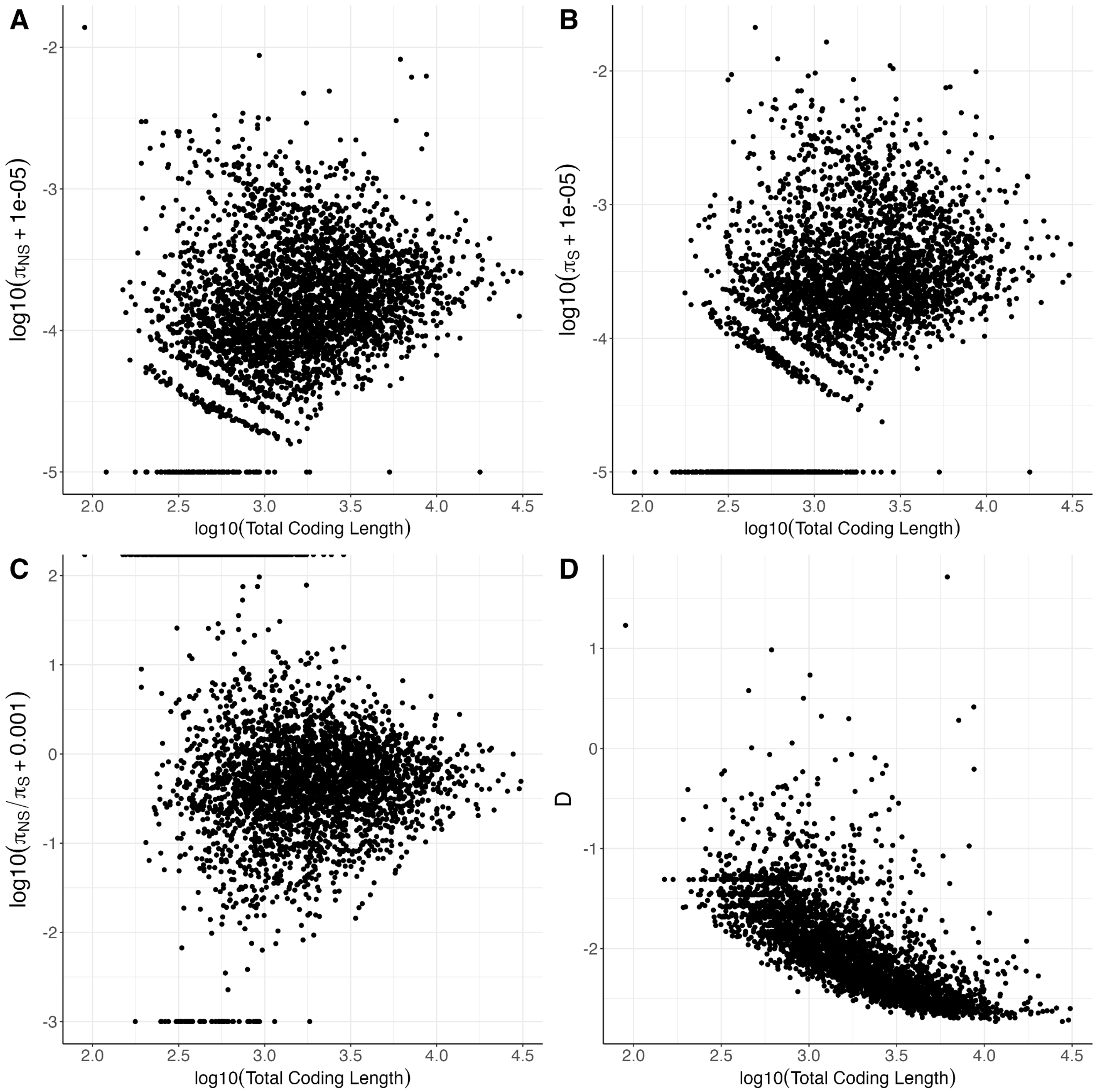


**Figure S5.** Genome-wide correlations of $\boldsymbol{\pi}_{\mathrm{NS}}$ (A), $\boldsymbol{\pi}_{\mathrm{S}}$ (B), $\boldsymbol{\pi}_{\mathrm{NS}}/\boldsymbol{\pi}_{\mathrm{S}}$ (C), and Tajima's *D* (D) with the log10-scaled total coding length of transcripts (Kendall's $\boldsymbol{\tau}$ = 0.190, 0.264, -0.0978, -0.605; $p < 2.2 \times 10^{-16}$, $p < 2.2 \times 10^{-16}$, $p < 2.2 \times 10^{-16}$, $p < 2.2 \times 10^{-16}$). To avoid excluding zeros in the visualizations, we incremented all values by the lowest observed order of magnitude for the statistic (A-C).

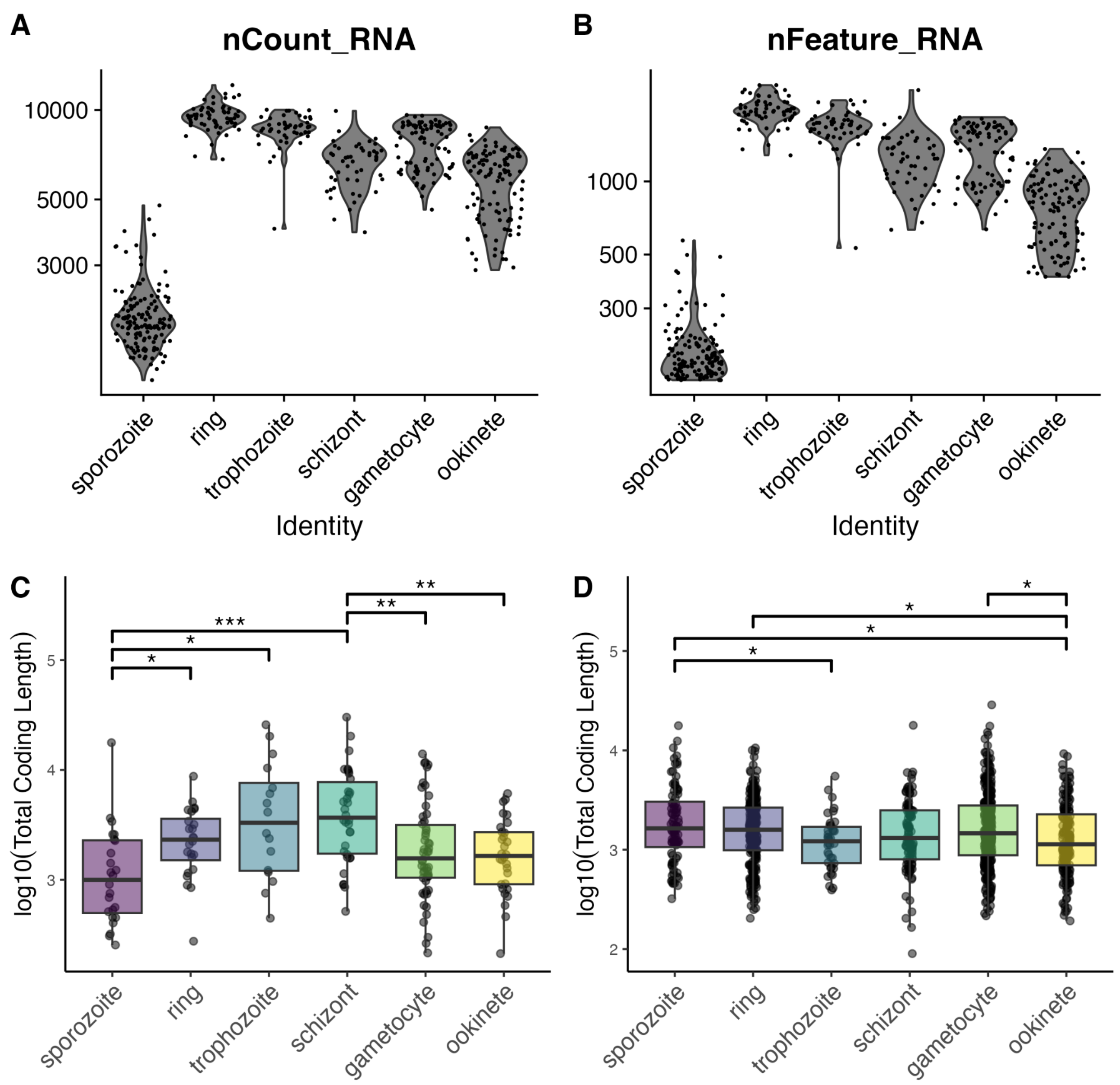


**Figure S6.** Assay-level differences in stage-specific features. After initial cell filtering steps (retaining cells with minimum 150 features and less than 5% mitochondrial and apicomplexan DNA), cell counts in Malaria Cell Atlas *P. falciparum* Smart-seq2 dataset assay varied by stage (sporozoite *N*=141, ring *N*=63, trophozoite *N*=51, schizont *N*=50, gametocyte *N*=88, ookinete *N*=112). Total (A) and unique (B) number of RNA transcripts by cell type annotation after cell- and transcript-level filtering. Coding length (log10 scale) of primary gene sets by life stage for (C) primary gene sets (BH-adjusted significant Pairwise Wilcoxon Rank Sum comparisons: sporozoite-ring, $p$ = 0.0166; sporozoite-trophozoite, $p$ = 0.0106; sporozoite-schizont, $p$ = 0.000461; gametocyte-schizont, $p$ = 0.00472; ookinete-schizont, $p$ = 0.00474) and (D) secondary gene sets (BH-adjusted significant Pairwise Wilcoxon Rank Sum comparisons: ookinete-sporozoite, $p$ = 0.0361, ookinete-ring, $p$ = 0.0361, ookinete-gametocyte, $p$ = 0.0437;

trophozoite-sporozoite, $p = 0.0493$). Significant comparisons in (C) and (D) between stages marked with asterisks (*$p < 0.05$, **$p < 0.01$, ***$p < 0.001$).

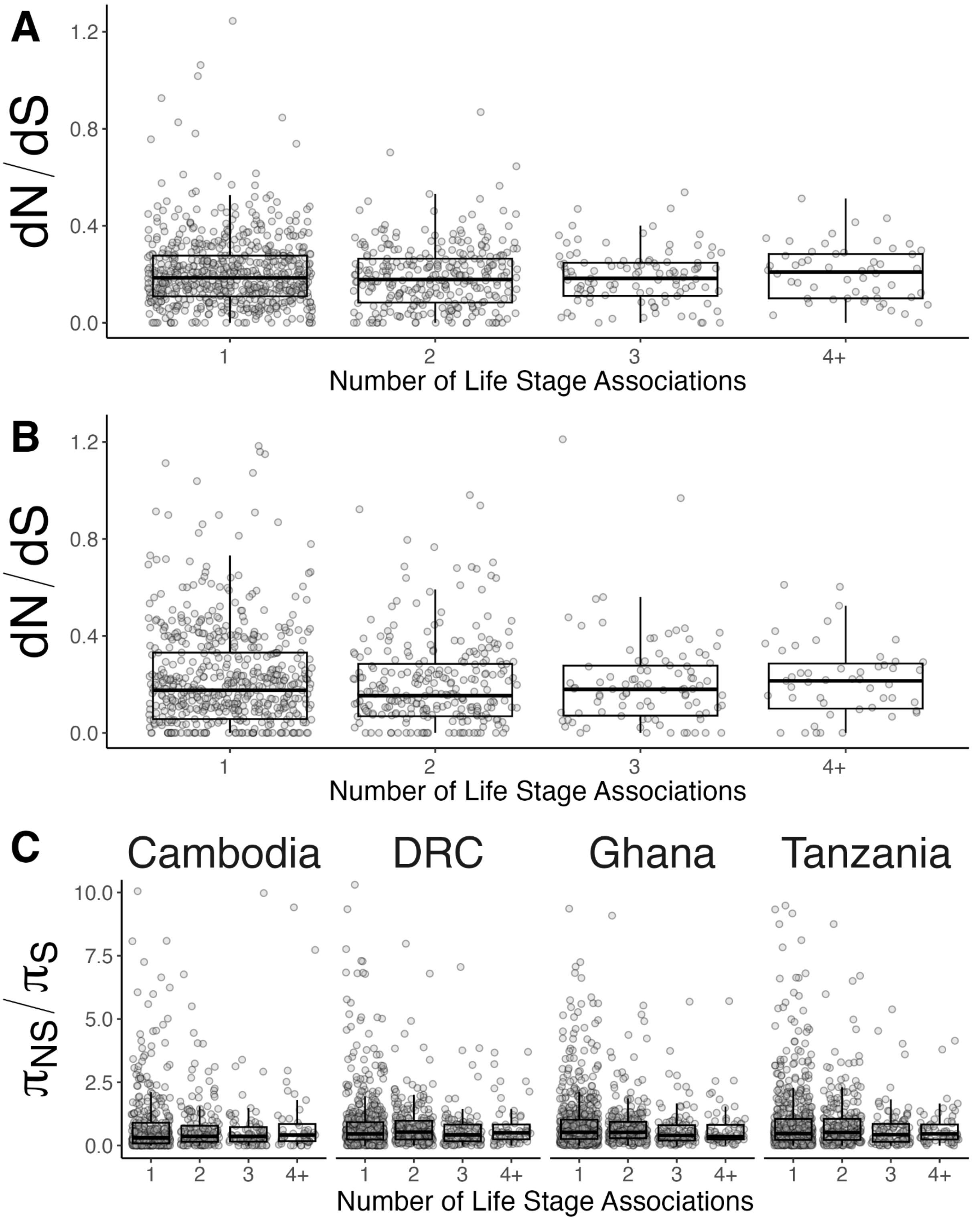


**Figure S7.** (A) *dN/dS* of *P. falciparum-P.reichenowi* orthologs by breadth of expression across the *P. falciparum* life cycle (Partial Spearman's $\rho$ = -0.112, $p$ = $1.32 \times 10^{-4}$) [3 values above y-axis limit not shown 3 infinite *dN/dS* values not represented]. (B) *dN/dS* of *P. falciparum-P. praefalciparum* orthologs by breadth of expression across the *P. falciparum* life cycle (Partial

Spearman's $\rho$ = -0.0969, $p$ = 0.00122) [5 values above y-axis limit not shown and 48 infinite *dN/dS* values not represented]. (C) $\pi_{NS}/\pi_{S}$ by breadth of expression across the *P. falciparum* life cycle in four distinct populations. In three of four populations, $\pi_{NS}/\pi_{S}$ trends toward a weak decrease with breadth (Cambodia $\rho$ = -0.00565, $p$ = 0.851; DRC $\rho$ = 0.00872, $p$ = 0.759; Ghana $\rho$ = -0.0439, $p$ = 0.120; Tanzania $\rho$ = -0.0202, p = 0.474) [19 outliers above y-axis limit and 798 infinite values not represented].

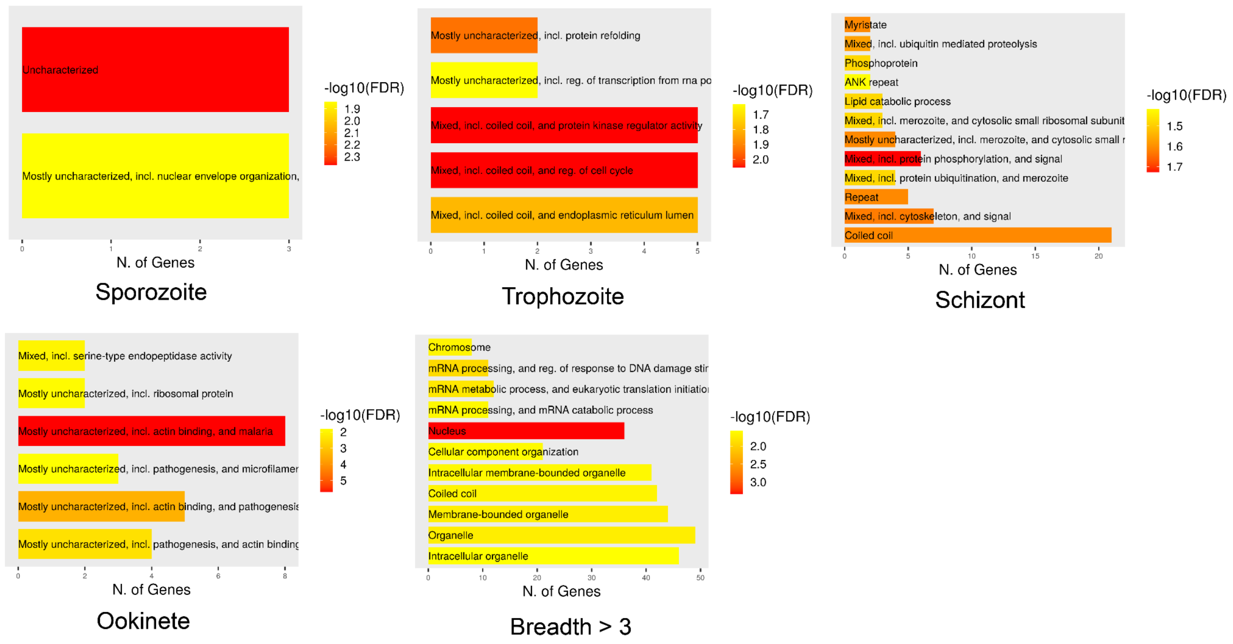


**Figure S8.** Functional enrichments for gene sets with single stage expression determined by approach 1 and for genes with breadth of expression above 3 stages, including the number of enriched genes with FDR < 0.05 (x-axis), function (y-axis/bar label), and -log10FDR (color). No significant functional enrichments were found for the ring and gametocyte gene sets. GSEA analyses were conducted and plotted in ShinyGO v. 0.81 utilizing the *Plasmodium falciparum* 3D7 STRINGdb dataset as the genomic background.

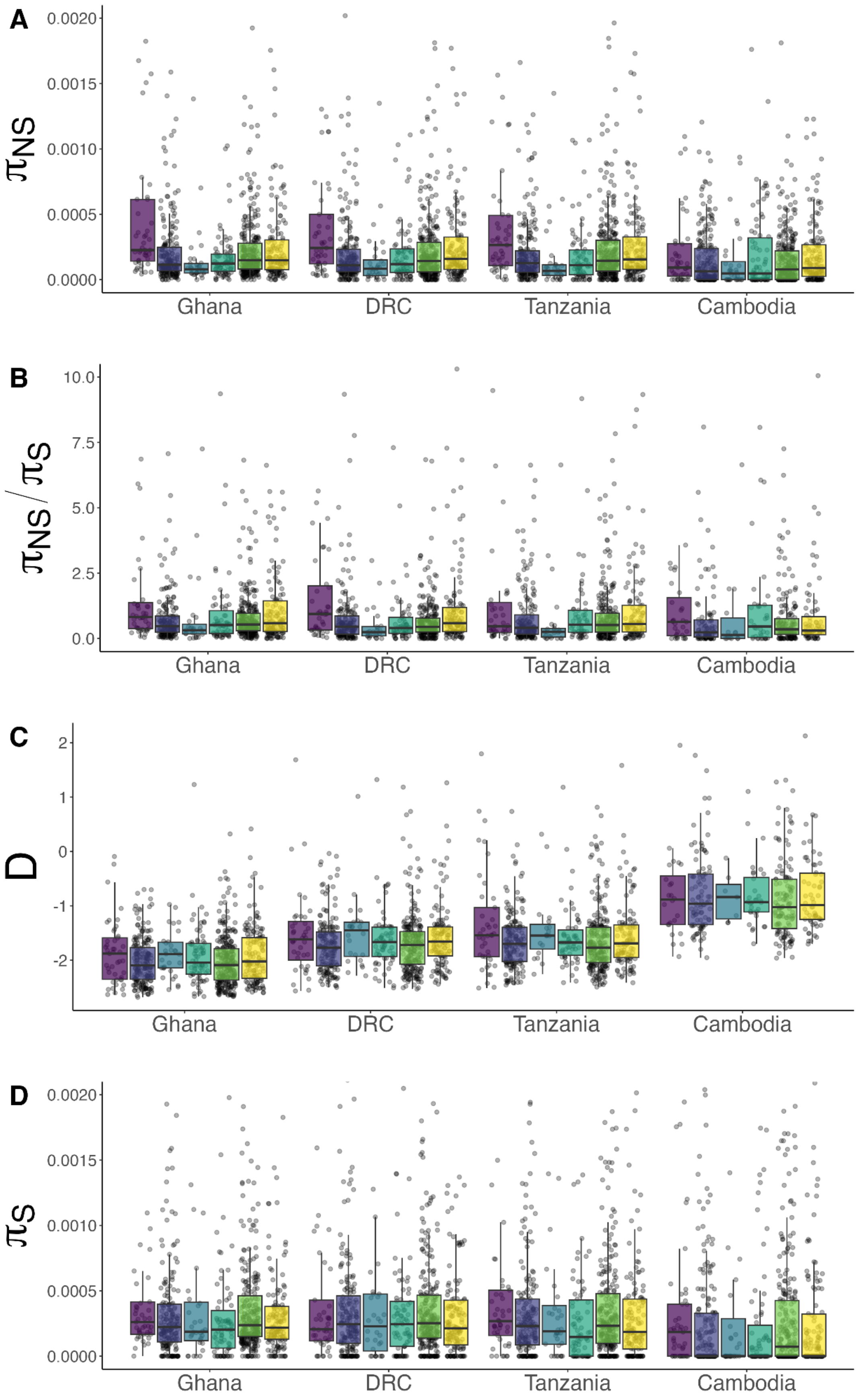

A
0.0020
0.0015
0.0010
0.0005
0.0000
$\pi_{NS}$
Ghana
DRC
Tanzania
Cambodia
B
10.0
7.5
5.0
2.5
0.0
$\pi_{NS}/\pi_S$
Ghana
DRC
Tanzania
Cambodia
C
2
1
0
-1
-2
D
Ghana
DRC
Tanzania
Cambodia
D
0.0020
0.0015
0.0010
0.0005
0.0000
$\pi_S$
Ghana
DRC
Tanzania
Cambodia

**Figure S9.** Gene-level population diversity statistics including $\boldsymbol{\pi}_{NS}$ [33 outliers above y-axis limit not shown] (A) $\boldsymbol{\pi}_{NS}/\boldsymbol{\pi}_{S}$ [18 outliers above y-axis limit not shown and 524 infinite values not represented] (B), Tajima's *D* (C), and pairwise $\boldsymbol{\pi}_{S}$ [144 outliers above y-axis limit not shown] (D) across four geographically diverse MalariaGen parasite population samples (x-axis). Estimates are shown for genes with single-stage expression and grouped by life stage, which is denoted by boxplot color (purple: sporozoite, periwinkle: ring, blue: trophozoite, blue-green: schizont, green: gametocyte, yellow: ookinete). Gene sets with expression in a single life stage used here are defined using the secondary approach relying on a combination of 50% overlap and differential expression criteria (Methods).

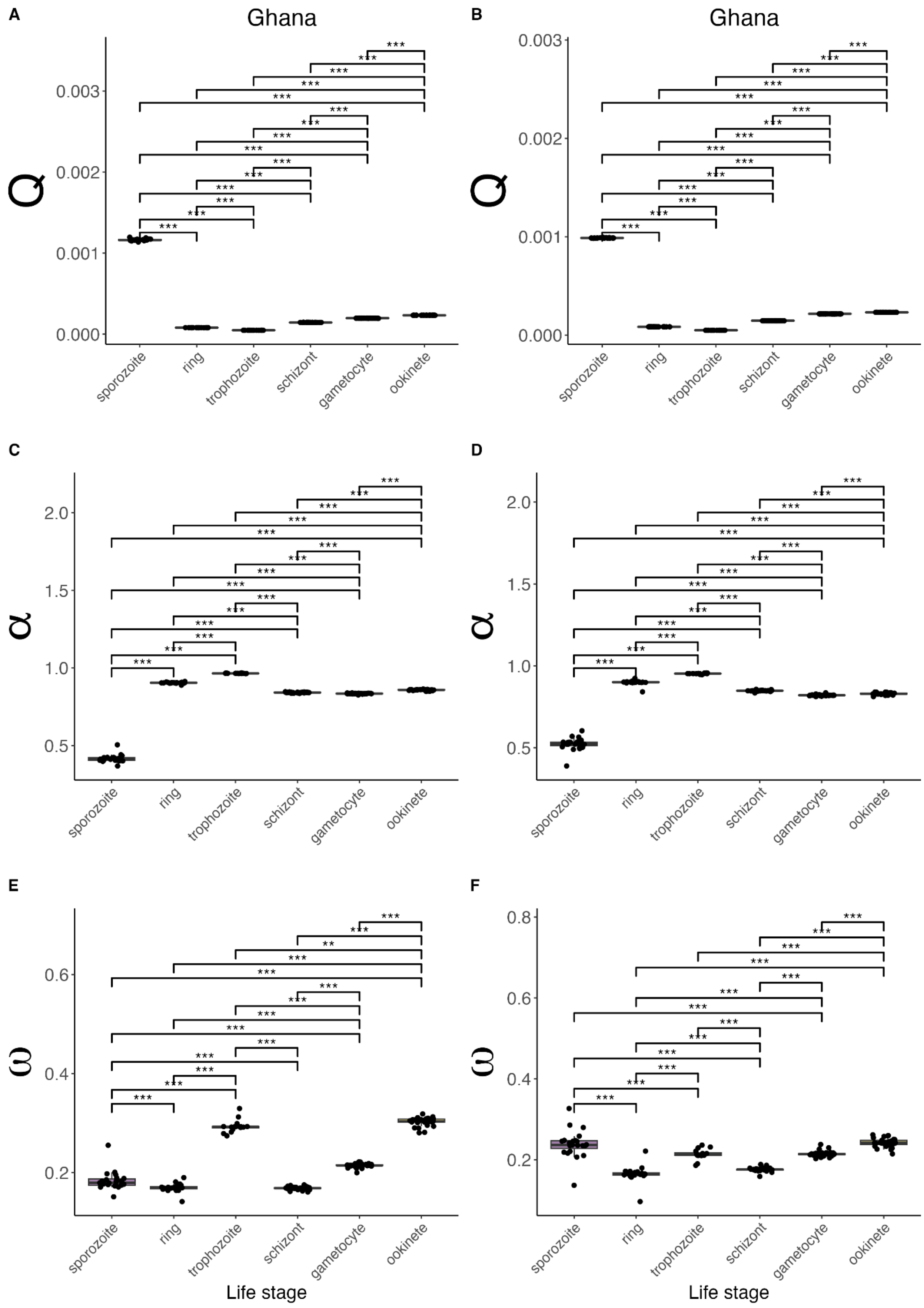
A
Ghana
B
Ghana
Q
0.000
0.001
0.002
0.003
C
D
α
0.5
1.0
1.5
2.0
E
F
ω
0.2
0.4
0.6
0.8
sporozoite
ring
trophozoite
schizont
gametocyte
ookinete
***
**
Life stage

**Figure S10.** Jackknifed estimates of probability of fixation of a deleterious mutation (*Q*) (A-B), Proportion of adaptive substitution (*α*) (C-D), and the relative rate of adaptive divergence (*ω*) (E-F) estimated by DFE-alpha analysis of site frequency spectra from the Pf7 parasite population in Ghana and *P. falciparum*-*P. reichenowi* (A, C, E) or *P. falciparum*-*P. praefalciparum* (B, D, F) divergence estimates for nonsynonymous (selected) and fourfold degenerate (A, C, E) or synonymous (B, D, F) (neutral) SNPs in *P. falciparum* life stage-associated genes. All stage-stage pairwise comparisons of jackknifed estimates revealed significant differences by life stage (Bonferroni-adjusted Pairwise Wilcoxon rank sum test ***$p < 0.001$).

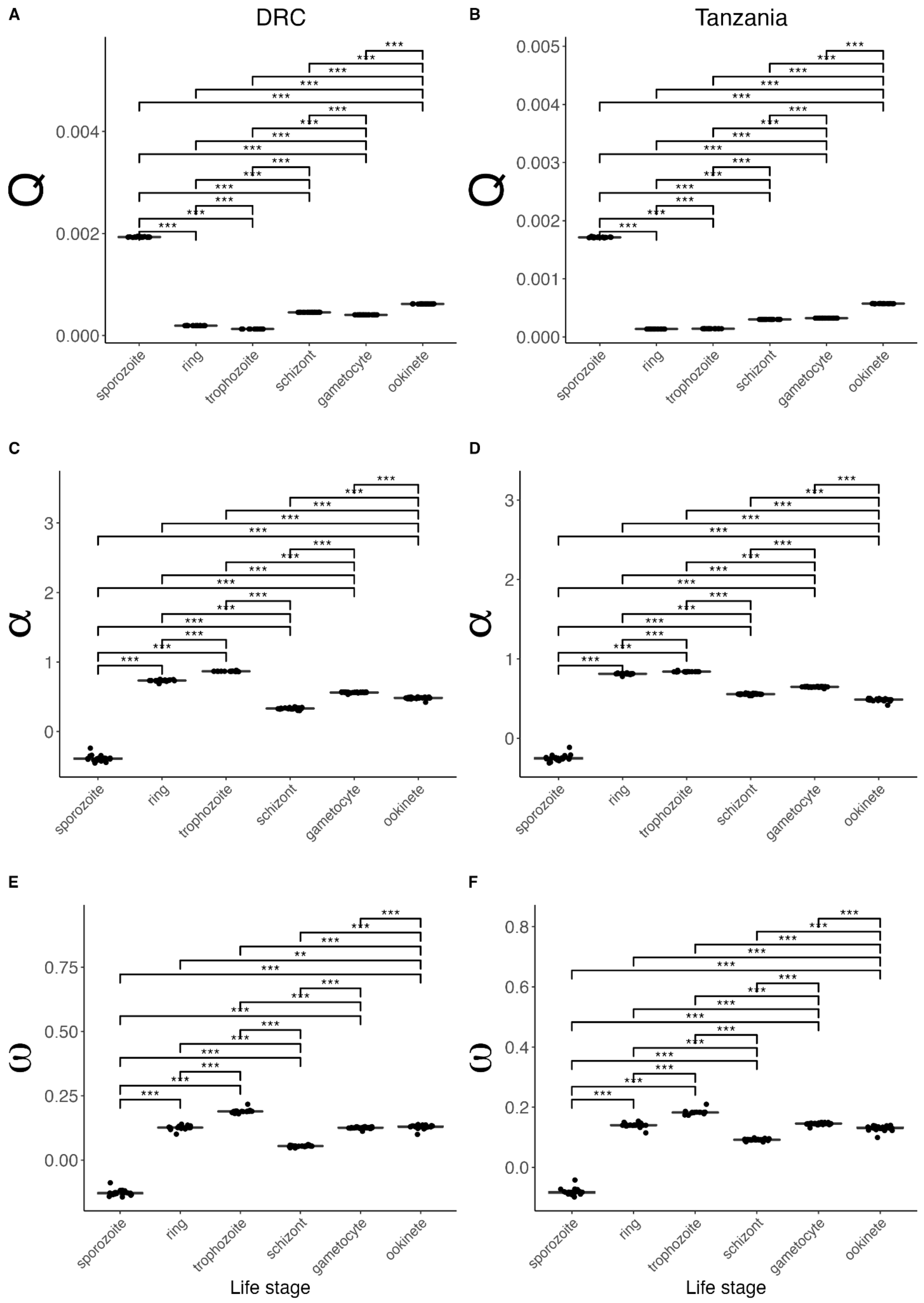

A
DRC
B
Tanzania
Q
0.000
0.002
0.004
0.001
0.003
0.005
C
D
α
0
1
2
3
E
F
ω
0.00
0.25
0.50
0.75
0.0
0.2
0.4
0.6
0.8
***
**
sporozoite
ring
trophozoite
schizont
gametocyte
ookinete
Life stage

**Figure S11.** Jackknifed estimates of probability of fixation of a deleterious mutation (*Q*) (A-B), Proportion of adaptive substitution ($\alpha$) (B), and the relative rate of adaptive divergence ($\omega$) (C) estimated by DFE-alpha analysis of site frequency spectra from different Pf7 populations (A,C,E: DRC, B,D,F: Tanzania) and *P. falciparum*-*P. reichenowi* divergence estimates for nonsynonymous (selected) and synonymous (neutral) SNPs in *P. falciparum* life stage-associated genes. All stage-stage pairwise comparisons of jackknifed estimates revealed significant differences by life stage (Bonferroni-adjusted Pairwise Wilcoxon rank sum test $^{**}p < 0.01$, $^{***}p < 0.001$).

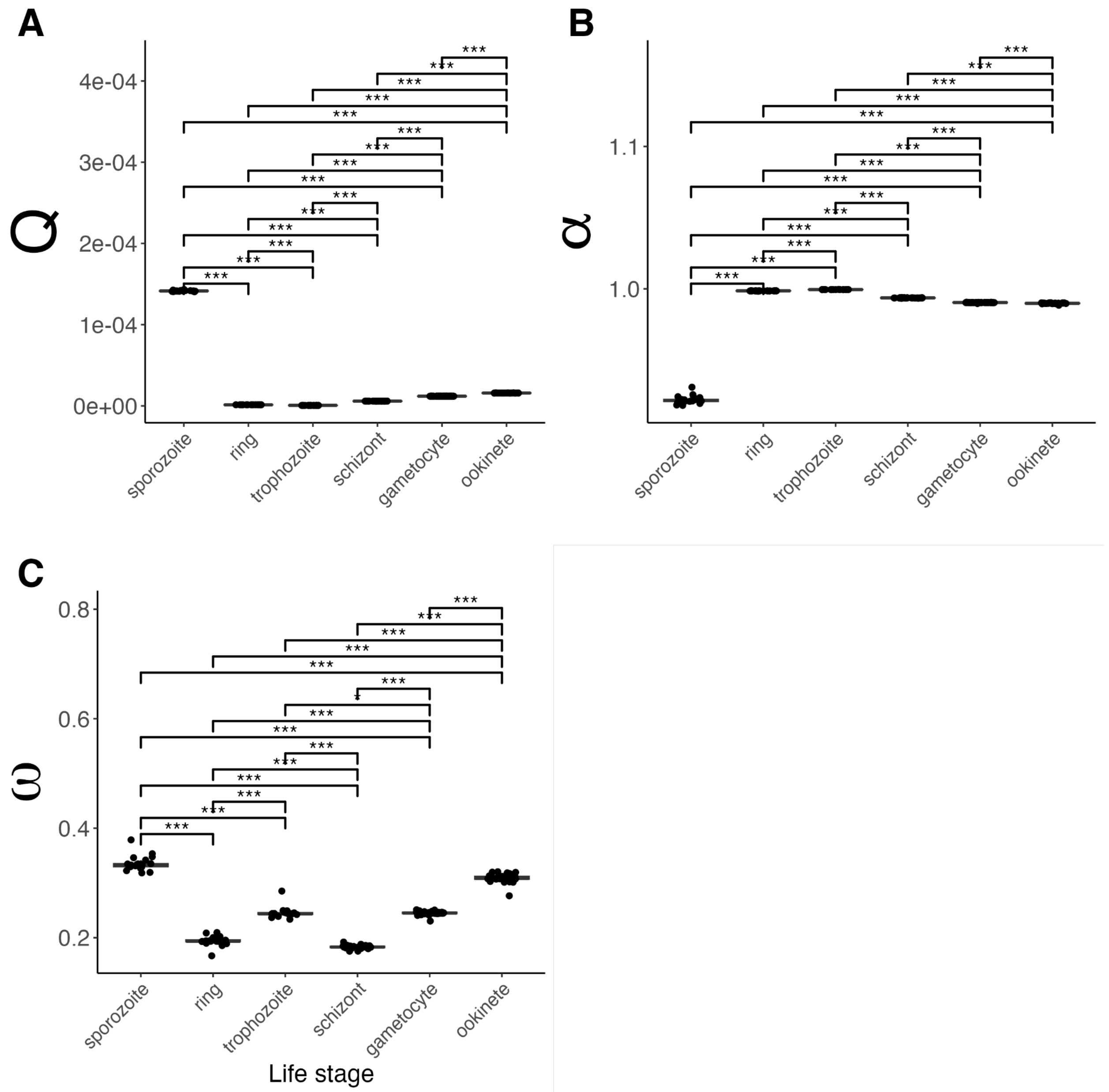


**Figure S12.** Jackknifed estimates of probability of fixation of a deleterious mutation (*Q*), Proportion of adaptive substitution (*α*), and the relative rate of adaptive divergence (*ω*) estimated by DFE-alpha analysis of site frequency spectra and *P. falciparum*-*P. reichenowi* divergence estimates for nonsynonymous (selected) and synonymous (neutral) SNPs in *P. falciparum* life stage-associated genes. This analysis does not use the demographic expansion model to correct for recent *P. falciparum* bottlenecks. All stage-stage pairwise comparisons of jackknifed estimates revealed significant differences by life stage (Bonferroni-adjusted Pairwise Wilcoxon rank sum test *$p < 0.05$, **$p < 0.01$, ***$p < 0.001$).

**Table S1.** FDR-corrected (BH) significance *Q*-values for meta-analysis of Tukey test differences in mean per-gene diversity statistics by stage (as defined in the primary gene sets) across four geographically diverse *P. falciparum* populations. FDR corrections were applied separately to each diversity statistic, reflecting the non-independence of the statistics shown.

| Pairwise stage-specific differences in diversity AWFisher q-values | | | | |
|---|---|---|---|---|
| Comparison | $\pi_{\mathbf{NS}}$ | $\pi_{\mathbf{S}}$ | $\frac{\pi_{\mathbf{NS}}}{\pi_{\mathbf{S}}}$ | $D$ |
| ring - sporozoite | **1.13e−4** | 0.997 | **4.42e−5** | **1.17e−6** |
| trophozoite - sporozoite | 0.856 | 0.997 | **0.0023** | **0.0309** |
| schizont - sporozoite | **9.66E − 5** | 0.997 | **1.70e−5** | **0.00111** |
| gametocyte - sporozoite | **3.17e−6** | 0.997 | **4.60e−4** | **1.17e−6** |
| ookinete - sporozoite | **0.00577** | **0.00543** | **0.00319** | 0.105 |
| trophozoite - ring | 0.997 | 0.997 | 0.947 | 0.719 |
| schizont - ring | 0.997 | 0.997 | 0.947 | 0.719 |
| gametocyte - ring | 0.997 | 0.997 | 0.947 | 0.719 |
| ookinete - ring | 0.997 | 0.0735 | 0.947 | **0.0309** |
| schizont - trophozoite | 0.997 | 0.997 | 0.947 | 0.923 |
| gametocyte - trophozoite | 0.997 | 0.997 | 0.947 | 0.775 |
| ookinete - trophozoite | 0.997 | 0.0572 | 0.947 | 0.775 |
| gametocyte - schizont | 0.997 | 0.997 | 0.947 | 0.775 |
| ookinete - schizont | 0.997 | 0.0735 | 0.947 | 0.775 |
| ookinete - gametocyte | 0.997 | **6.68e−4** | 0.947 | 0.707 |

**Table S2.** FDR-corrected (BH) significance *Q*-values for meta-analysis of Tukey test differences in mean per-gene diversity statistics by stage (as defined in the alternative gene sets) across four geographically diverse *P. falciparum* populations. FDR corrections were applied separately to each diversity statistic, reflecting the non-independence of the statistics shown.

| Pairwise stage-specific differences in diversity AWFisher q-values | | | | |
|---|---|---|---|---|
| Comparison | $\pi_{\mathbf{NS}}$ | $\pi_{\mathbf{S}}$ | $\frac{\pi_{\mathbf{NS}}}{\pi_{\mathbf{S}}}$ | $D$ |
| ring - sporozoite | **9.10e−6** | 0.996 | **2.68e−4** | **0.0141** |
| trophozoite - sporozoite | **7.16e−4** | 0.996 | **2.11e−5** | 0.912 |
| schizont - sporozoite | 0.186 | 0.996 | **0.00344** | 0.33 |
| gametocyte - sporozoite | **3.44e−5** | 0.996 | **0.00284** | **0.00239** |
| ookinete - sporozoite | 0.0818 | 0.996 | 0.201 | 0.0672 |
| trophozoite - ring | 0.954 | 0.996 | 0.183 | 0.984 |
| schizont - ring | 0.228 | 0.996 | 0.692 | 0.984 |
| gametocyte - ring | 0.954 | 0.996 | 0.263 | 0.984 |
| ookinete - ring | 0.0818 | 0.996 | **2.68e−4** | 0.984 |
| schizont - trophozoite | 0.321 | 0.996 | 0.0979 | 0.984 |
| gametocyte - trophozoite | 0.945 | 0.996 | **0.0114** | 0.984 |
| ookinete - trophozoite | 0.369 | 0.996 | **1.34e−4** | 0.984 |
| gametocyte - schizont | 0.494 | 0.996 | 0.930 | 0.984 |
| ookinete - schizont | 0.861 | 0.996 | **0.0297** | 0.984 |
| ookinete - gametocyte | 0.0818 | 0.996 | **0.0114** | 0.984 |

**Table S3.** FDR-corrected (BH) *Q*-values for meta-analysis of Tukey test differences in mean per-gene Hudson F-statistics by stage (as defined in the primary gene sets) across all pairwise comparisons of four *P. falciparum* populations.

| Pairwise stage-specific differences in $F_{ST}$ AWFisher q-values | |
|---|---|
| Comparison | $\mathbf{F_{ST}}$ |
| ring - sporozoite | **2.59e−4** |
| trophozoite - sporozoite | **0.0144** |
| schizont - sporozoite | 0.0907 |
| gametocyte - sporozoite | **2.59e−4** |
| ookinete - sporozoite | **0.0132** |
| trophozoite - ring | 0.875 |
| schizont - ring | 0.875 |
| gametocyte - ring | 0.883 |
| ookinete - ring | 0.672 |
| schizont - trophozoite | 0.883 |
| gametocyte - trophozoite | 0.875 |
| ookinete - trophozoite | 0.875 |
| gametocyte - schizont | 0.855 |
| ookinete - schizont | 0.883 |
| ookinete - gametocyte | 0.875 |
